# Opening the black box: a modular approach to spike sorting

**DOI:** 10.64898/2026.01.23.701239

**Authors:** Samuel Garcia, Chris Halcrow, Charlie Windolf, Zachary M. McKenzie, Paul Adkisson-Floro, Heberto Ramon Mayorquin, Benjamin Dichter, Alessio P. Buccino, Pierre Yger

## Abstract

Spike sorting is an algorithmic process that extracts the activity of individual neurons from extracellular electrophysiology recordings. With the ballooning use of high density probes, such as Neuropixels, this essential processing step is increasingly becoming time consuming and computationally expensive. Although many software tools have been proposed to address spike sorting, they are usually constructed and benchmarked as monolithic “black boxes”, making it difficult to factor out the effects of individual algorithmic steps on the final outcome, especially when varying datasets and parameters. To address this issue, we developed a modular and common framework to develop, benchmark and assemble the key computational steps that are used in state-of-the-art spike sorting algorithms. Relying on fast and efficient ground truth generation of biophysically plausible recordings, we show that we are able to individually benchmark and precisely quantify the performance of different steps in a spike sorting pipeline (i.e. peak detection, feature extraction and clustering, and template matching). We then leverage these results to create a modular, component-based spike sorter that can outperform Kilosort 4 on dense and large simulated recordings and produce similar quantitative results on real data. In addition, we find that the major bottleneck of all modern spike sorting pipelines is in the physical motion of probes, regardless of the drift-correction strategy. The component-based spike sorting framework presented here has the potential to foster community engagement in the field by lowering the barrier to contributions and providing a flexible yet powerful framework to construct end-to-end spike sorting solutions.

## Introduction

Spike sorting is a processing step to extract individual spike times of individual neurons from extracellular recordings [1, 2]. With the development and widespread adoption of high-density micro-electrode arrays both for *in-vivo* [3, 4, 5] and *in-vitro* applications [6, 7, 8], automated solutions to the spike sorting problem are essential. In recent years, the neuroscience community has therefore produced a wide range of open-source tools to optimize spike sorting [9, 10, 11, 12, 13, 14, 15, 16, 17, 18].

Almost all spike sorting algorithms follow a similar data processing pipeline which we break down into five components: preprocessing, peak detection, feature extraction, clustering, and template matching. In more detail: raw signals are preprocessed (sometimes corrected for probe motion), then a selection of putative spikes are detected. The features of these spikes are computed and used to form clusters. Templates, representing the spatial-temporal footprint of the “average” spike in the cluster, are then computed to redetect spikes using a deconvolution method. Different spike sorters use different methods for each sorting component (see Figure 1 comparing `Kilosort2`, `Kilosort4`, and `SpyKING-CIRCUS`). Despite this shared structure, each spike sorting tool is designed and used as an end-to-end “monolithic” pipeline, which leads to several problems.

**Figure 1:**
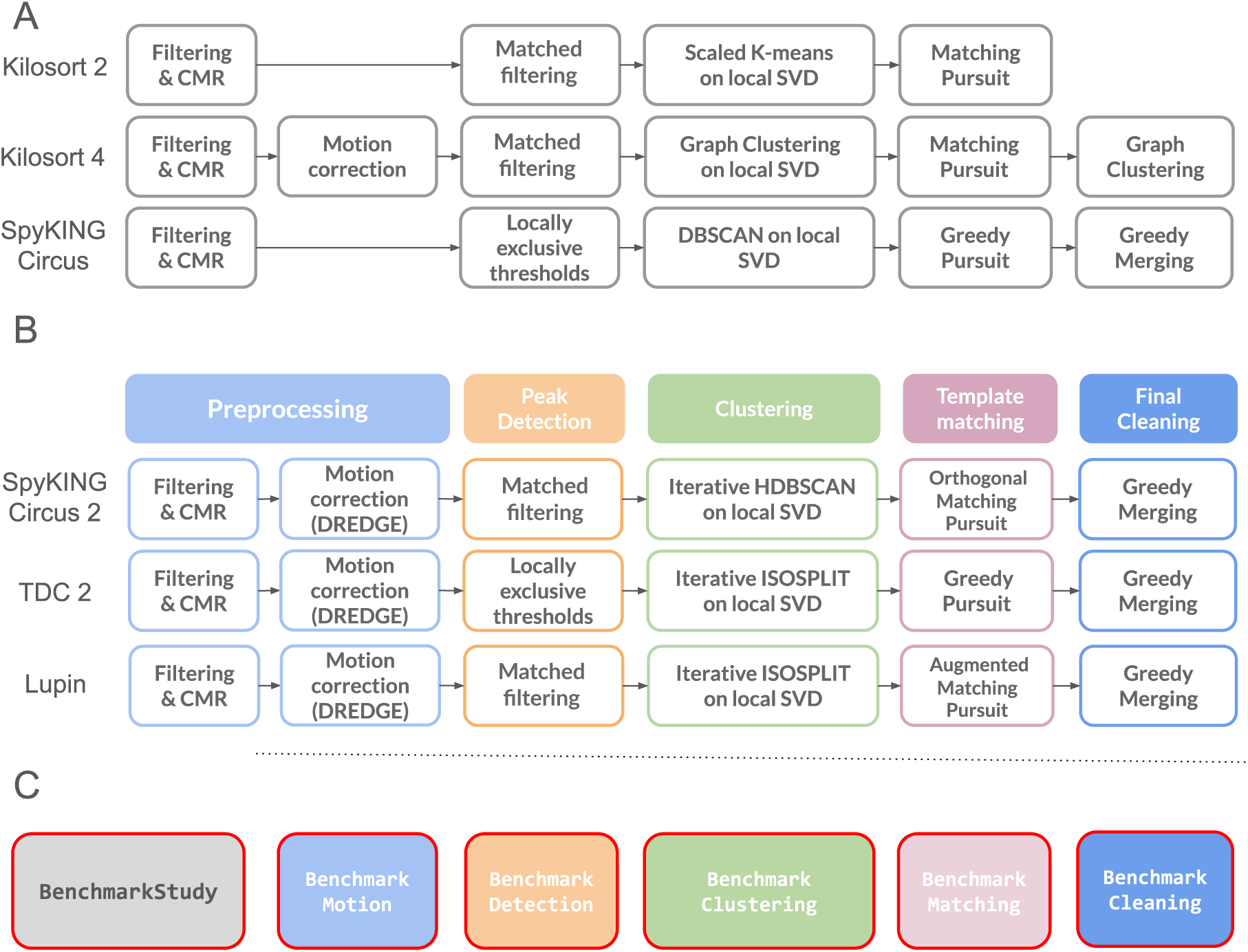
**A** Most of modern spike sorters tend to have the same sequence of algorithmic steps, with particular details and/or implementations choices. Particular examples are shown for Kilosort2 and 4 [18, 10], alongside SpyKING-CIRCUS [15]. **B** For each of these key algorithmic steps, we have factorized and re-implemented various algorithms to properly benchmark each of them on a per step basis, bypassing the need to compute performance on the whole sequence. **C** Relying on the modular architecture, BenchMarkStudy objects on a per step basis can be created, to test/optimize individual components.

First, it is difficult to test, implement, and contribute new methods for individual spike sorting steps. Although there has been continuous algorithmic development in spike sorting with new ideas and concepts used to tackle motion correction [19, 20, 10], peak detection [21, 22, 23], feature extraction and clustering [24, 25, 26, 27, 28], and template matching [29, 30, 31], these potentially powerful methods for spike sorting remain largely unused, mainly because they are not embedded in a ready-to-use spike sorting tool. This challenge creates a very high-barrier for developers to enter the spike sorting field.

Second, it is difficult to benchmark new algorithmic ideas. Although notable efforts towards end-to-end spike sorting comparisons have been proposed [32], direct benchmarks of individual spike sorting steps are still limited and scattered in the field. Every new contribution, such as a new peak detection or clustering method, requires a large overhead effort to compare it to other methods, i.e., by using ad-hoc and non-validated re-implementations, or the development of sub-optimal integrated pipelines to compare with other existing tools. It is also currently very difficult to ask questions of the type “how does a new peak detection method affect the downstream results of a given spike sorter?”.

Finally, it is a challenge for developers to maintain their spike sorting packages. The maintenance burden of the tools falls almost exclusively on the original developers, who, in most cases, follow academic incentives [33] and move on to other projects. In an era where hardware and software are in constant development, software maintenance is an essential part of software development. For example, Python versions have an official support of 5 years from their release (Python 3.10, released in late 2021, will reach end-of-life in October 2026), making software developed in older Python versions virtually unmaintainable. Similarly, MATLAB versions have rolling support for GPU hardware, which makes GPU-accelerated software developed in older MATLAB versions (e.g., Kilosort 1/2/2.5/3) difficult to sync with modern hardware. In general, software maintenance problems can render otherwise state-of-the-art tools unusable within a few years of their development. This is a genuine problem in academic software development, and most of the spike sorting tools mentioned above are currently unmaintained^1^: Klusta [9], JRClust/IronClust [13], SpyKING-CIRCUS [15], HDSort [14], WaveClus [16], YASS [17].

In this paper, we propose an alternative framework to provide a modular and community-based paradigm for spike sorting development. Building on the SpikeInterface framework [34], a mature and well-maintained software ecosystem for extracellular electrophysiology data analysis, we introduce a new sortingcomponents module. This module provides a flexible, modular framework based on the five sorting components discussed earlier. The sortingcomponents module includes several methods for each component, with an extensible framework to support new contributions. Each component is paired with a benchmarking tool, allowing for easy comparisons between existing methods and new ones. We believe that the approach implemented here can solve many of the issues detailed above.

To kickstart the proposed community effort, we have re-implemented or ported several methods available in the literature and from existing spike sorting tools. In the following sections, we first introduce the sortingcomponents modular framework and present a fast and efficient ground-truth simulation module, which includes drift simulation [19]. Next, we showcase the framework with a detailed benchmark of the currently available methods for peak detection, feature extraction plus clustering, and template matching. We then go into detail on a template matching failure mode, leveraging our ground-truth simulations to pinpoint issues with interpolation as a source of spike sorting failures for recordings with drift. Finally, we compare end-to-end spike sorting solutions implemented *via* this sortingcomponents-based solution with Kilosort4 [18], which is arguably the most widely used state-of-the-art spike sorter in the field. To demonstrate the net gain of using this component-based approach, we present three new sorting pipelines built with the modular approach of the components. The first two originally motivated the development of such a modular architecture: SpyKING-CIRCUS 2, an enhanced version of SpyKING-CIRCUS [15], and TriDesClous 2. The third, Lupin, is a new component-based sorter created as the optimal combination of the best-performing components, based on our generated data and benchmarks. In our simulated benchmarks, and on real-world datasets, we show that Lupin is competitive with Kilosort4 both in performance and speed, highlighting the benefit of step-wise benchmarks and modularity.

## Results

### A modular approach to the spike sorting problem

In recent years, a broad consensus has emerged among the latest spike sorting algorithms regarding how a typical algorithmic pipeline should look. On a macroscopic scale, and taking into account some internal subtleties in preprocessing, most modern algorithms are now structured like those shown in Figure 1A. This involves preprocessing (typically filtering and denoising *via* Common Median Reference [CMR] and whitening), peak detection, feature extraction, then clustering to obtain a dictionary of templates, followed by a template matching step and a final gathering and/or cleaning of the results. Although the exact nature of all these steps may vary and is beyond the scope of this current paper (see Methods of the aforementioned spike sorters), all modern spike sorters, including `Kilosort` [10, 18], `SpyKING-CIRCUS` [15], `MountainSort` [12], `YASS` [17] and `TriDesClous` follow such a pipeline. However, these pipelines are always evaluated as “black boxes” [32] so that it is hard to properly assess the pros and cons of each of the individual steps.

We have implemented a new sortingcomponents module in the popular SpikeInterface package. We call each step in a spike sorter a component, and one can now implement a full spike sorter as a series of components, adjusting each one for fine granularity over their pipeline. As suggested in Figure 1B, we port three new spike sorters to this conceptual architecture: SpyKING-CIRCUS 2 and TriDesClous 2 are direct evolutions of previous algorithms, while Lupin is a new spike sorting algorithm (see Methods), created to demonstrate the power of such modularity. Although the three pipelines share similar processing pipelines, sometimes even using similar algorithms, they differ by the particular parameters of the algorithms used, and thus each have different pros and cons, as will be shown in this paper.

Crucially, each key algorithmic step is paired with a higher-order object termed a BenchmarkStudy, see Figure 1C, enabling performance benchmarking at the component level for each algorithm. The exact nature of each benchmark depends on the component, e.g., how many peaks are detected for a detection method or how many clusters can be found for a clustering method. The benchmarks allow us to identify strengths and weaknesses of each step in spike sorting pipelines.

The sortingcomponents module uses a built-in engine to parallelize the computational load, which splits the recordings into temporal chunks. This ensures that the flexibility and modularity offered do not adversely affect the processing times of the algorithms. The BenchmarkStudy object can be used to compare different methods in a given step or to compare different parameters of the same method. This new object, integrating the built-in comparison features of SpikeInterface, can reduce the time needed for developers to implement a new algorithmic idea and compare it to other available methods without benchmarking the whole pipeline, which can itself bias the results. This modular framework thus turns spike sorting into a community-based effort, where the pros and cons of individual steps can be efficiently understood, and the best ones reused in many different pipelines.

### A fast and efficient ground truth generation module

In order to quickly benchmark the numerous components of the spike sorting pipelines, one needs to be able to generate artificial ground truth data in a fast and efficient manner. We have designed and implemented a new way to generate artificial ground truth recordings within SpikeInterface. As shown in Figure 2A, ground truth recordings with any number of neurons can be generated on-the-fly once a probe layout is provided. Each neuron can have a unique spatio-temporal waveform (also known as a template), which can be generated from a mathematical model or loaded from external libraries, as well as a position and firing rate. Each probe channel can have its own noise profile, with or without correlations between channels.

**Figure 2:**
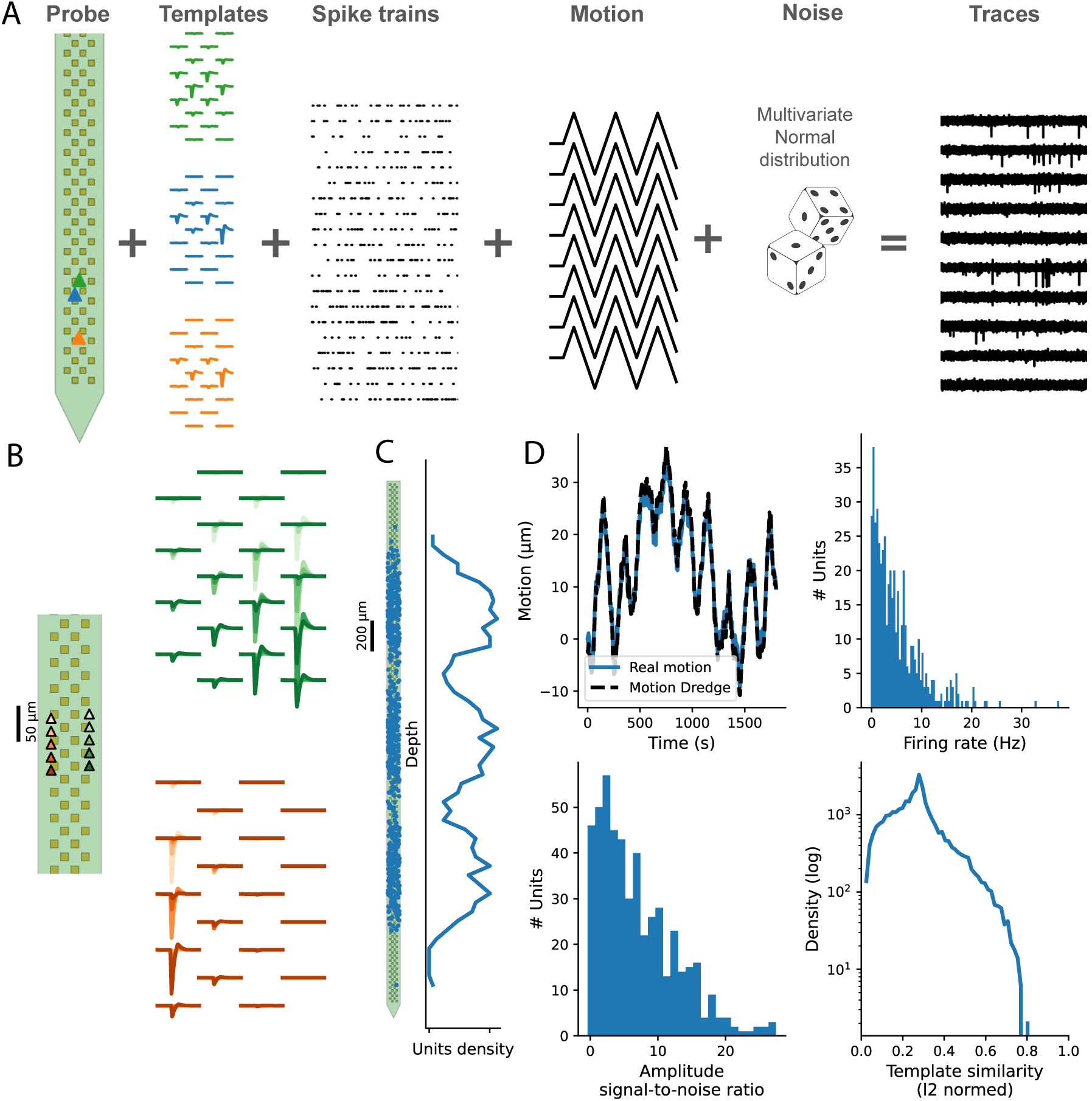
Ground truth recording generation. **A** Given a probe layout, templates, activity patterns and inhomogeneous motion vectors, we can generate, on-demand and lazily, extracellular traces to extensively benchmark spike sorting. **B** Left: Positions of two neurons (triangles) during drift, color-coded as a function of position. Right: During drift extra-cellular templates will be dynamically generated as functions of positions of the sources. Each line display the extracellular voltage on a particular spatial channel, colored to match the positions on the left plot. **C** The positions of the cells are drawn randomly from a trimodal Gaussian along the depth of a Neuropixels-like probe (see Methods) **D** Top left: the rigid motion vector impacting all the cells during the course of the recording. Bottom Left: the distribution of the signal-to-noise ratio for all the templates generated given the cells positions. Top Right: the distribution of firing rates for all the cells in the recording. Bottom Right: the distribution of Euclidean distances between the templates, for all pairs of cells in the recording, on a log scale.

To challenge modern spike sorters and make our ground truth recordings more realistic, we allowed for the inclusion of possible motion (drift) of the probe. As noted in recent papers [1], this is currently one of the major bottlenecks in spike sorting, but the algorithmic methods designed to solve the problem are not entirely satisfactory [19]. In our generator, neurons can drift with precomputed and imposed motion vectors, allowing for non-uniform motion that changes as a function of the neuronal depth along the recording probes. If neuronal waveforms are generated from a mathematical model (see Methods), as drift occurs the waveforms are modified accordingly as functions of the positions of the somas (see Figure 2B). If templates are taken from external libraries, the templates are interpolated as functions of the drift. In our generator, we model noise as a multivariate normal distribution, but other options could be implemented (1/f and/or multi-unit activity such as in [18], many background units [35], or accurate noise spectrum [36]).

In the rest of the paper, we will use 30 minute long recordings with a Neuropixels 1.0-like layout (384 channels), sampled at 30 kHz, with a trimodal distribution profile for the positions of the cells (see Figure 2C). In Figure 2D, we show some macroscopic statistics from an example artificially generated recording (see Methods for more details). The rigid motion was generated as a random walk, while firing rates were drawn from a gamma distribution (see Methods). The analytical model used to generate the templates as a function of the position provides a large diversity of template shapes and amplitudes. This is visible in the distribution of signal-to-noise ratios for the templates and in the distribution of Euclidean distances between pairs of templates.

### Benchmarking individual spike sorting steps improves overall spike sorting quality

To demonstrate the power of our modular approach to spike sorting, we combined the ground truth generator with the BenchmarkStudy objects to assess where mistakes in spike sorting algorithms originate.

#### Peak detection

The first step we assess with the components framework is peak detection. In peak detection we aim to identify all putative spiking events in a recording. Here, we compare the two most commonly used peak detection methods from recent literature: “locally exclusive” and “matched filtering” (see Methods). The first method detects peaks as local extrema within a spatio-temporal exclusion radius (to take into account the fact that when the channel density is high, action potentials emitted by neurons are likely to be seen on many channels simultaneously). The second method, introduced in [4, 18], relies on the concept of matched filtering [37]. This method can boost the signal-to-noise ratio of the extracellular signal, assuming we approximately know the spatio-temporal shapes of the motifs we are looking for.

To examine the difference between these two peak detection algorithms, we generated one static ground truth recording (see Methods) and assessed how many ground truth peaks each method detected. Figure 3 A displays the distribution of ground truth peaks as a function of peak amplitude, and the distribution of peaks that each method detected. We see that the locally exclusive method^2^, finds very few peaks below the detection threshold; whereas the matched filtering method can recover some peaks below threshold. The locally exclusive method seems to be better at finding very large (< −`200`µV) peaks. This could be partially explained by spike collisions: when two large spikes overlap in space and time, the resulting waveform could be highly distorted. Since matched filtering looks for “stereotypical” shapes, it might miss these events. The locally exclusive method could still detect these events since it simply looks at threshold crossings.

**Figure 3:**
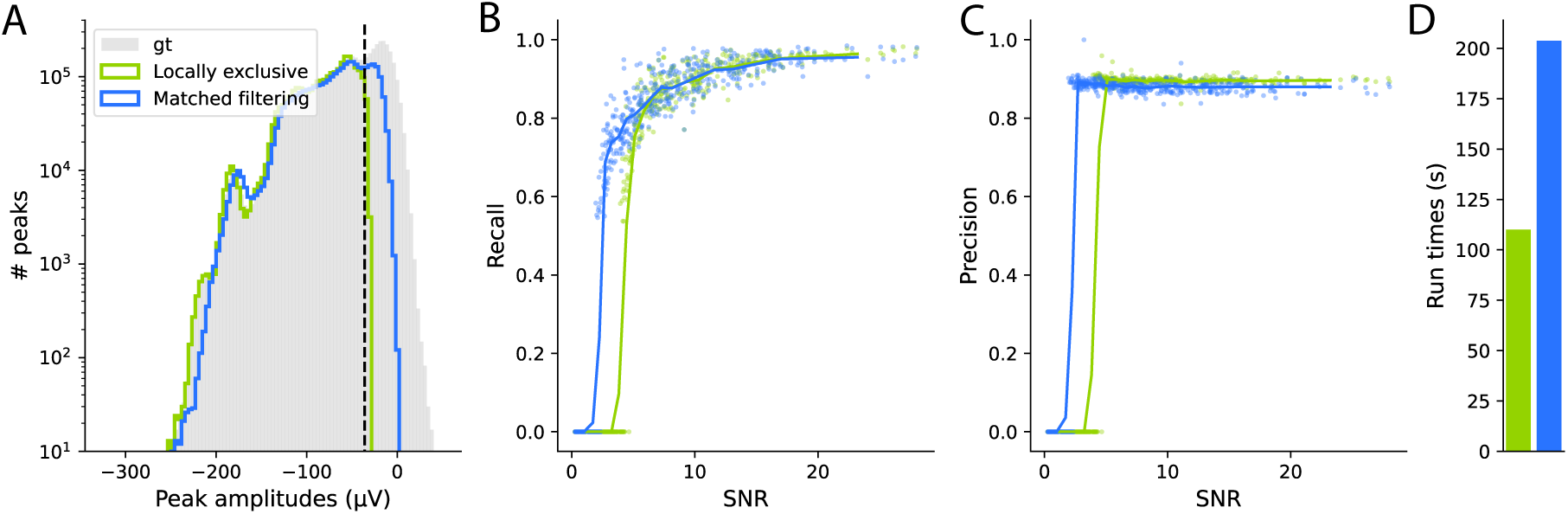
Peak detection benchmark. **A** Given the true distribution of the peak amplitudes in the ground truth recordings (gray area), we compute the recovered distribution for the peaks detected either via the “locally exclusive” method (blue), or “matched filtering” (green). **B** The recall for each neuron shown as dots in the ground truth recording as function of their signal-to-noise ratios for the two aforementioned methods. Lines show the binned average SNR **C** Same as in **B**, but for Precision. **D** The run times (in seconds) for the two methods.

In Figure 3 B and C we plot the recall and precision for each unit from a single generated recording for both methods. These are plotted as functions of the signal-to-noise ratio (SNR) of the units with a rolling average overlaid. The plots show that for low SNR, recall and precision are higher for the matched filtering method. Even with matched filtering, the recall starts to decrease for SNRs lower than 20. This means that even for quite large peaks relative to the noise, some are missed because of collisions and/or noise. This observation is a strong argument for additional steps, such as template matching, to recover these putative spikes. Note that matched filtering should be most beneficial when the electrode density is high, as it uses the spatial redundancy of spike detectability. In Figure 3 D we plot the run times of the two methods. We see that, although matched filtered has performed better overall, especially with small peaks, this performance comes with some computational cost.

In modern spike sorters, the peaks computed in this step are not used directly as the final spike times of neurons. Instead, they are used to build the clusters and templates of each unit. The final spike times are computed during the later template matching step, which has been proven to outperform other methods [39]. Hence, it is not clear if missing spikes during peak detection has a large impact on overall spike sorting.

#### Feature extraction and clustering

The next component we evaluated is clustering. In this step, each algorithm receives all spiking events as input and clusters them into groups representing putative units. It is a crucial, if not the most crucial, step in spike sorting to disambiguate individual neurons. Before clustering, we must extract features from each event. Because most spike sorters currently rely on Singular Value Decomposition (see [2] for a review) for feature extraction, e.g. [40, 15, 17, 41], we decided to only focus on benchmarking the clustering step. Therefore, in the following, we will consider the feature extraction and the clustering as a whole rather than individually.

In order to quantify the effects of motion correction on each clustering algorithm, we applied our analysis to two conceptually similar datasets: one termed “static” and one termed “motion-corrected”. Both datasets have the same initial cell input. For the motion-corrected recording, we imposed rigid motion on this input, then used `DREDge` to estimate the motion and finally interpolated the recording using kriging (see Methods and [19, 20]). Hence, if the motion was perfectly computed and compensated for, the static and motion-corrected recordings would be equivalent.

In Figure 4A (top), we compare four clustering algorithms that we have extracted and re-implemented from various spike sorting tools. All these algorithms are given the same inputs (i.e. the exact peaks times from the ground-truth recordings) and are evaluated in a similar manner, suppressing any biases that might be due to differences in peak detection methods. Given the fact that we are providing the theoretically ideal input data to these algorithms, the ground truth spike times, we expect to get an upper bound of their performances.

**Figure 4:**
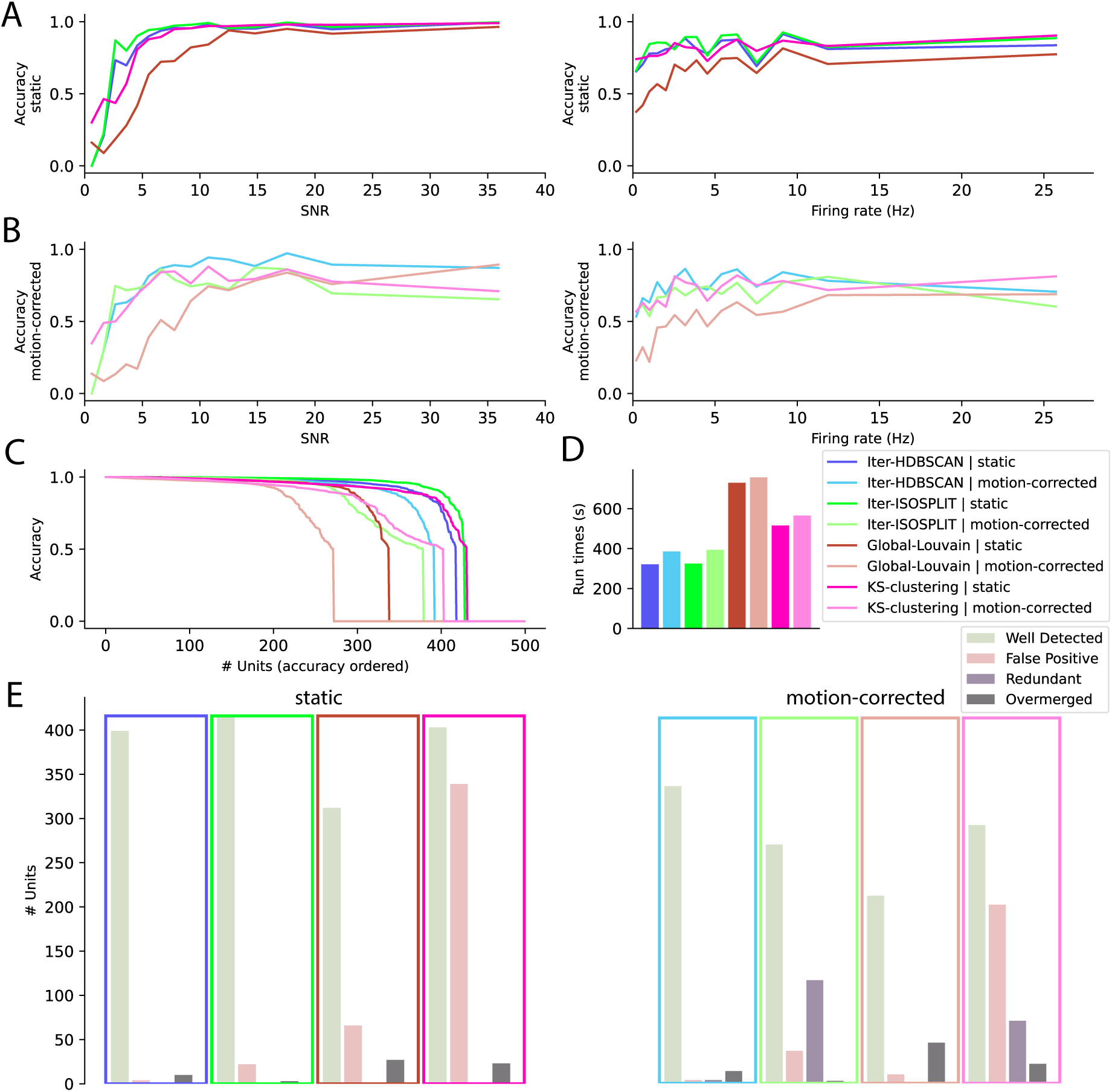
Feature extraction and clustering benchmark. **A** Accuracy recall for all the clustering methods when applied on true peak times from a static recording. Left: as a function of the signal-to-noise ratios of the neurons. Right: as a function of the firing-rate of the neurons **B** Same as in **A**, but applied to motion-corrected recordings, with the exact same activity/firing as the static one (see Methods) **C** Sorted accuracy levels for all clustering methods and recording types, as function of the neurons present in the artificial recordings **D** The run times for all the methods and datasets **E** For all clustering methods and recording types, the number of Well Detected, False Positive, Redundant and Overmerged units (see Methods).

Figure 4A compares the performance of several algorithms (see Methods for a description of their implementations). All methods perform well on static data. Iter-HDBSCAN (used in SpyKING-CIRCUS 2 [15]) and Iter-ISOSPLIT (used in TriDesClous 2 and Lupin) are both based on iterative splits of local density-based clustering performed by grouping the spikes per electrode, and give comparable results. KS-clustering is a re-implementation of the local graph clustering found in [18]. Global-Louvain is an attempt to design a single unified graph-based clustering on sparse connectivity matrices.

Interestingly we found, in Figure 4C, the number of units detected by the sorters is already far from perfect on static recordings (at best, 420 out of 500 units). We observe that clustering fails mainly for small SNR units, as opposed to small firing rate units (see Figure 4A left vs right). Hence, the failure to observe all units is driven by units with small SNRs being indistinguishable from noise.

The performance of all clustering algorithms drastically degrades for motion-corrected data. As seen in Figure 4B and C the clustering algorithms struggle to properly find the units, in contrast to the static case, suggesting that the motion-correction algorithms do not completely compensate for the probe motion. We can see a dramatic drop in performance, specifically in the number of units outside of the well-detected units category (see Figure 4E), for methods that rely on local clustering (such as KS-clustering). Here again, the influence of the signal to noise ratio seems to prevail over firing rates (see Figure 4B, right). Regarding the run times (Figure 4D), it is important to stress that Iterative methods of the sortingcomponents framework (Iter-HDBSCAN and Iter-ISOSPLIT) are fast as they rely heavily on parallelization over CPU-cores.

#### Template matching

We next assessed how effective the various template matching strategies used by different spike sorters are. In template matching we try to describe the incoming trace as a linear combination of spike waveforms, generated from a catalog of templates, plus noise. The temporal position of each waveform reveals the spike times while the template used from the catalog tells us the cluster index. In this way template matching simultaneously detects and clusters spikes. We compared the main algorithms available in the field, providing the ground truth template “catalog” as input. Although sharing similar principles, the exact mathematical nature of these algorithms is different. In the following, we compared the classical matching pursuit algorithm implemented in `Kilosort` (KS-matching [10]), a refined Orthogonal Template matching (Circus-OMP [42]), an augmented matching pursuit algorithm with temporal super-resolution (Wobble [17]), and a simpler greedy pursuit (TDC-peeler) (see Methods for more details of all these algorithms).

As seen in Figure 5A-C, all algorithms perform similarly at reconstructing the signal through template matching based approaches, with an accuracy that rises sharply as a function of the SNR of neurons, at least for static recordings (panel A). Methods based on full convolution of the traces with the catalog of templates (KS-matching, Circus-OMP and Wobble) have a better accuracy for small units compared to the greedy pursuit algorithm (TDC-peeler). However, once again, the performance for all algorithms is severely impacted for motion-corrected recordings (panel B; see Methods), since the interpolation of the traces considerably blurs the signals. Although there are some differences with respect to run times (Figure 5D), the user’s choice of template matching engine depends on some other factors (e.g. correlation levels, firing rates, etc.) that could be tested to make a well-informed choice depending on the input data. For example, Figure 5E shows the recall over a distribution of lags between spikes. Here, the Wobble algorithm has the best performance for spike collisions, i.e, spikes that overlap in space and time (the results are displayed for static recordings, but similar results are obtained for motion-corrected ones). Assuming that fine correlations are important to understand how information is finely encoded by a neuronal population, one might favor Augmented Matching Pursuit algorithms such as Wobble, even if they are slightly slower than the others.

**Figure 5:**
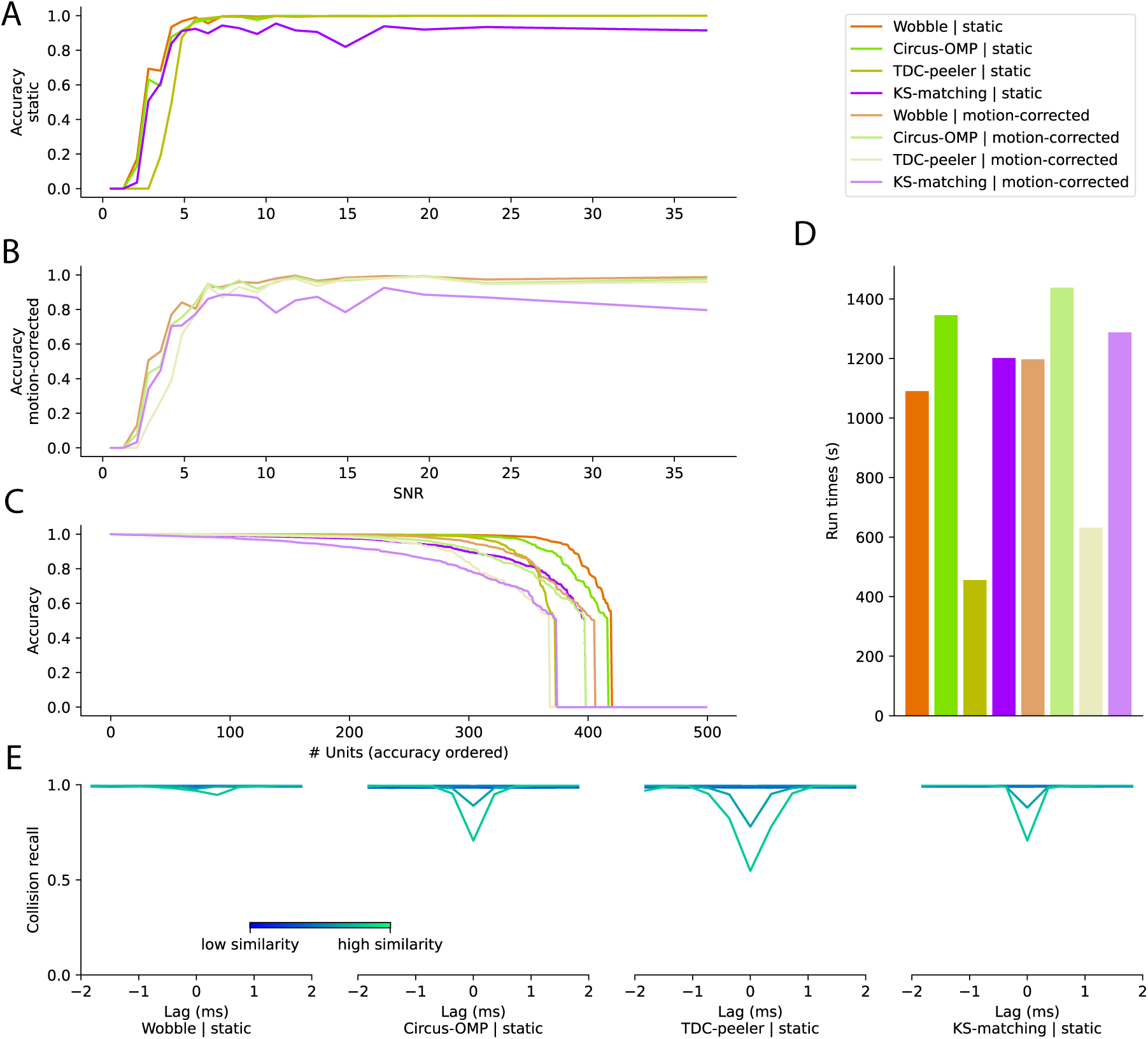
Template matching benchmark. **A** Accuracy recall for all the matching methods when launched with a perfect catalog of templates, and as a function of the signal-to-noise ratios of the neurons **B** Same as in **A**, but when applied to a motion-corrected recording, with the exact same activity/firing as the static one (see Methods) **C** Sorted accuracy levels for all matching methods and recording types, as a function of the neurons present in the artificial recordings **D** The run times for all the methods and datasets **E** For all matching methods, collisions levels [39] computed on the static recording, as a function of the temporal lag between all pairs of spikes, and the cosine similarity between pairs of templates.

### Motion interpolation reduces spike sorting performance

As shown in Figure 5, all template matching algorithms suffer from a lack of performance when dealing with motion-corrected recordings. This is not due to the estimation step, since motion can be estimated very accurately by the DREDge algorithm [20]. This spike sorting degradation occurs even when the ground truth motion is given. [19]. Instead, the performance drop is mainly due to the interpolation of the traces using the kriging kernel method [18]. This method constructs a kernel for every time bin of the estimated motion (1 s in our case) based on the motion vector, which is applied using a scalar product to the traces to compensate for motion. In short, every sample will be interpolated in space by the weighted average of the neighboring channels, which should compensate for the motion itself. Our previous work [19] has already shown a strong degradation in the performance of spike sorters when drift is present, which is mainly due to this interpolation step. Thanks to the modular approach introduced in this paper, we can now highlight how this performance loss affects clustering (Figure 4C-E) and template-matching (Figure 5C).

Knowing that motion interpolation causes a significant performance decrease in spike sorting, as shown in Figures 4C-5C, we implemented a new idea for template matching. Instead of interpolating the traces, we directly interpolated the templates [43]. This has three advantages: 1) the interpolation of the templates is less noisy because the templates are smoother than traces; 2) the templates can be pre-computed for some predefined motion steps (i.e., spatial bins) to cover the entire motion vector, resulting in faster computation; 3) the interpolation can be made via better methods (e.g. cubic interpolation). Because these interpolation methods are more computationally demanding they cannot be performed on-the-fly on raw traces, but can be performed on the templates. In Figure 6, we tested this idea of interpolating the templates instead of the traces during the template matching step. The core nature of the matching problem can be observed in Figure 6A. Here, we compare the “True” drifting templates (as defined by the generative mathematical model as a function of the cell position, see Methods) and the best interpolation that can be done with bicubic splines on a grid for a given motion vector. As can be seen, there are still some important non-zero residuals that give rise to errors.

**Figure 6:**
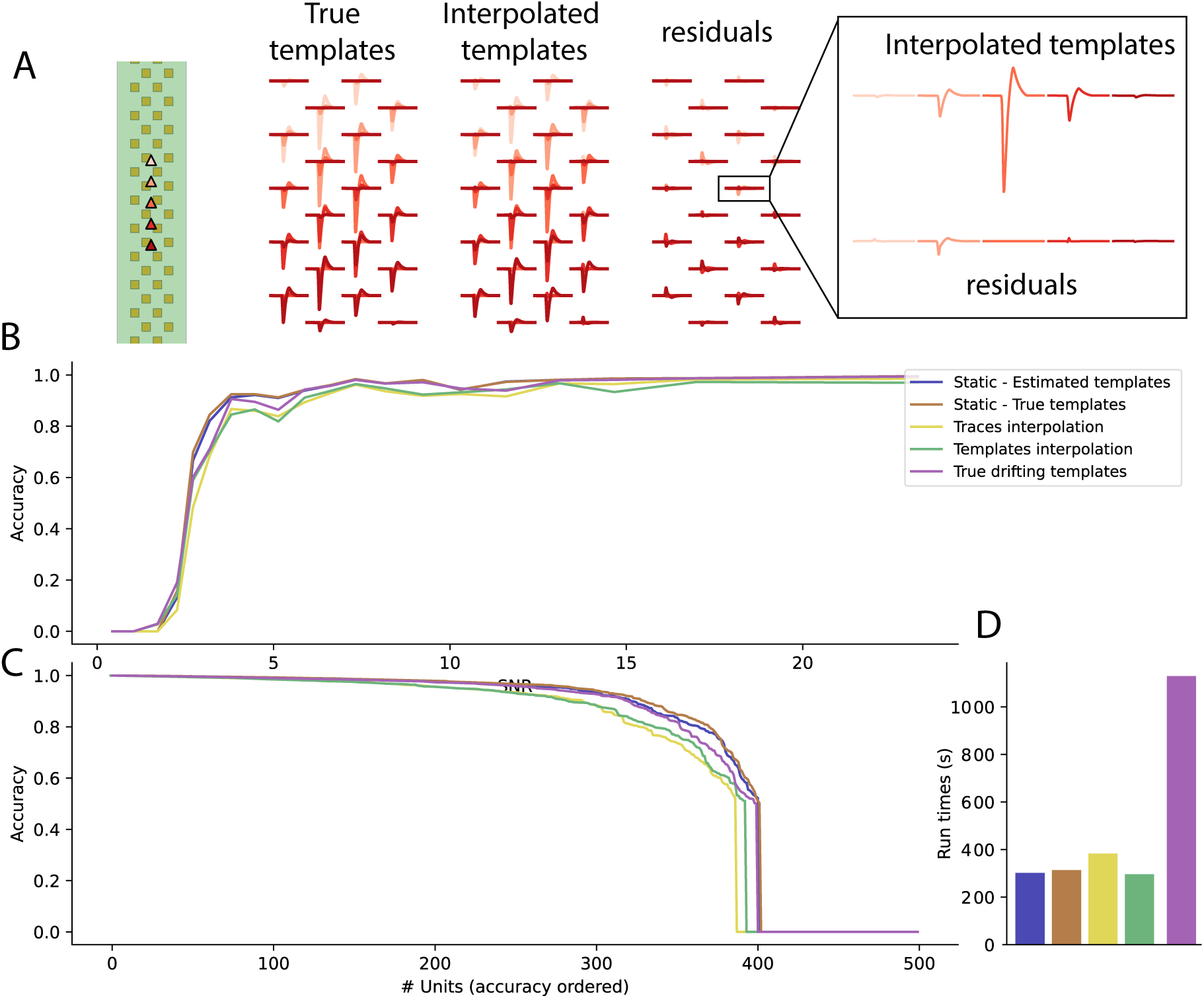
Motion correction strategies benchmark. **A** Example of template interpolation for a particular neuron drifting along the depth of the electrode. On the Left, the True templates are created from the generative mathematical model, taking into account the real position of the neuron. In the middle, templates are interpolated *via* a cubic spline interpolation for a particular displacement. On the right, the residuals (i.e. the differences) between estimated and real templates, as functions of the positions. The inset displays the interpolated templates and residuals on a single chosen channel, in a non-overlapping manner and as function of depth **B** Accuracy for several matching methods when run on a static recording, when Templates or Traces are interpolated given the estimated motion, or when we use the True static or drifting templates (perfect dictionaries of templates) and as function of the signal-to-noise ratios of the neurons **C** Sorted accuracy levels for all matching methods and types of dictionary, as function of the neurons present in the artificial recordings **D** The run times for all the methods and datasets.

To check the quantitative impact of these errors in the interpolation, we extended the Greedy Matching Pursuit algorithm (TDC-peeler) to have two modes: the standard one (where templates are kept fixed) and a new one with a “drifting templates" based algorithm, where we pre-compute interpolated versions of each template in a spatial range ([-100 µm, 100 µm]) for a given spatial step (1 µm)). In this new algorithm, instead of interpolating the traces, we interpolated templates at any given time and motion prior to the template-matching step. We compared these algorithms for template matching in five different situations:

1. “Static - Estimated Templates”: no drift is present in the recording, templates are estimated from raw data
2. “Static - True templates”: no drift is present, true templates are taken from the biophysical model (upper bound on static performance)
3. “Traces interpolation”: drift is present in the recording, traces are motion-corrected as a preprocessing step
4. “Templates interpolation”: drift is present in the recording, templates are estimated and then motion-corrected via bicubic splines before template matching
5. “True drifting templates”: drift is present in the recording, perfect templates from the biophysical model are used (upper bound on drifting performance)

The results are shown in Figure 6. As expected, the best performance occurs for static recordings. For recordings with drift, the performances of both traces-based and templates-based interpolations are similar and the accuracy is almost the same (Figure 6B,C). The results also show that estimating the templates from raw data or using real ground truth templates from the biophysical model has no clear impact on the performances for static recordings (Figure 6B,C). This is a good sign that, even for low-firing neurons, estimating the templates from a small subset of spikes does not dramatically degrade the matching procedure.

The final case “True drifting templates", although biologically impossible, provides an upper bound on the performance we could expect for matching in a drifting recording and allows us to estimate the cost of spatial interpolation to compensate for the drift. As shown in Figure 6B-C, the result is as we hypothesized: the matching performance of “True drifting templates” is not as good as for the static recordings, but is better than the other methods applied to drifting recordings (with an increased computational cost, Figure 6D, since true drifting templates are denser than interpolated ones). This leads us to the conclusion that, in a recording with drift, estimating the drift can be done accurately, but correcting for it through interpolation on traces or templates still greatly degrades the performance of spike sorting. This conclusion paves the way for future improvements, i.e. finding better ways of interpolating either templates or traces when motion is known or estimated.

### End-to-end evaluation of component-based sorters

Finally, the modular benchmark proposed in this article allows one to perform exhaustive comparisons of fully integrated spike sorters. In order to get a robust estimate of the performances, we computed the performance of five end-to-end spike sorters on multiple instances of artificially generated recordings, all with the same properties (see Methods). To demonstrate the full potential of the modular framework described in this paper, we compared a state-of-the-art spike sorting algorithm (`Kilosort4` [18]) to several new sorters entirely built on the components described in this article. We first developed `SpyKING-CIRCUS 2` and `TriDesClous 2`, as updated ports of `SpyKING-CIRCUS` and `TriDesClous`, to show that it was possible to create end-to-end components based sorters. We then constructed `Lupin` as the “best" sorter, based on the benchmarks in this paper. Based on Figures 3, 4 and 5 we choose matched filtering for peak detection, iterative ISOSPLIT for clustering and wobble (Augmented Matching Pursuit) for template matching. These three algorithms were originally developed as part of `Kilosort` [18], `Mountainsort` [12] and `YASS` [17] respectively, showing that `Lupin` benefits from ideas from an array of other spike sorters.

As can be seen in Figure 7A, the performances for all spike sorters on static recordings are comparatively strong, although all sorters sometimes miss obvious units with a large signal-to-noise ratio. This could be due to similar looking units, i.e., units whose physical positions are close in space, thus leading to similar extracellular waveforms that are hard to disambiguate for the clustering algorithms. Lupin is able to outperform all other spike sorters in both the static and motion-corrected cases.

**Figure 7:**
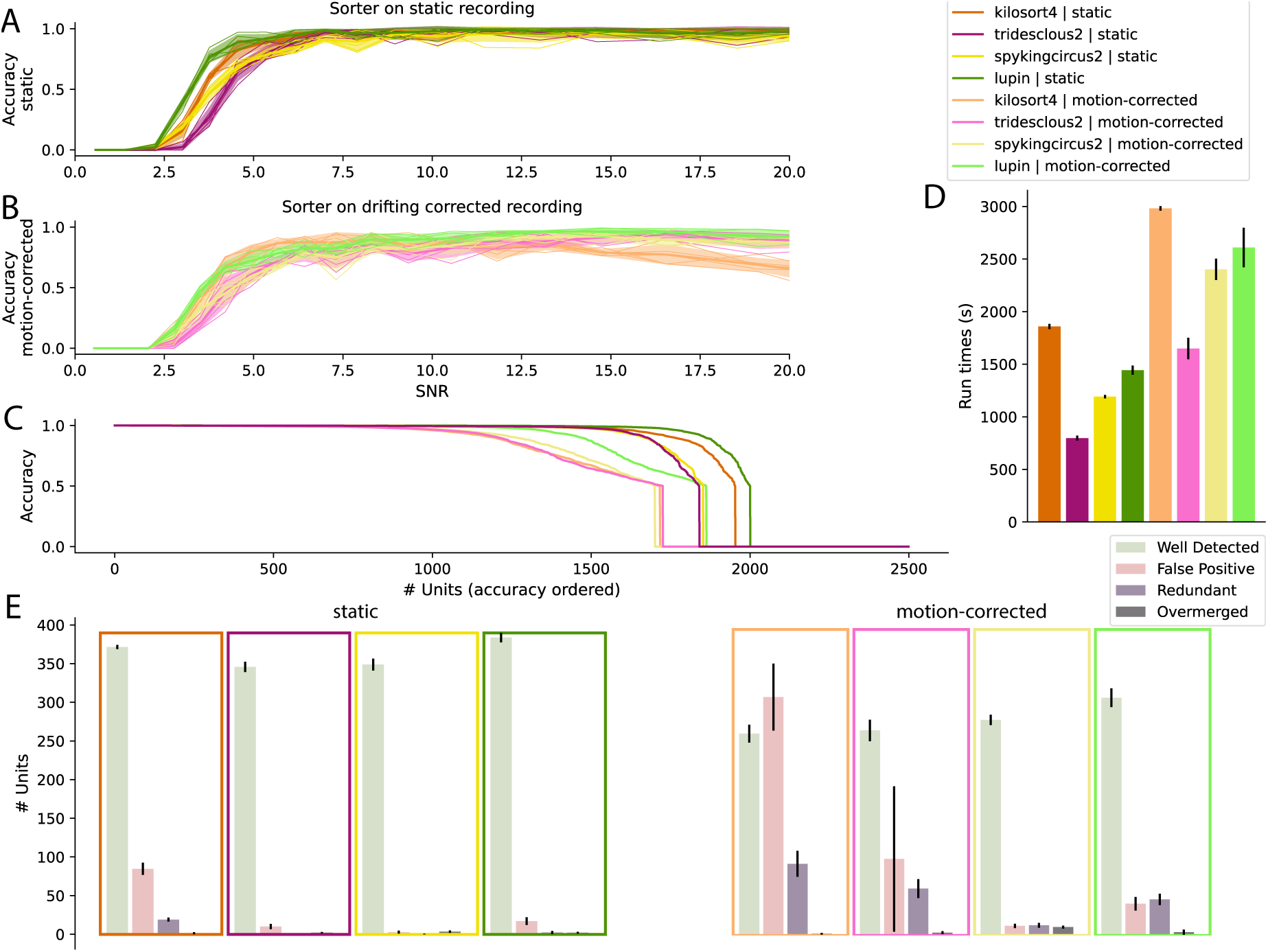
End-to-end spike sorter benchmark with simulated recordings. **A** Averaged accuracy for several spike sorters when applied to several static recordings (with various seeds), as function of the signal-to-noise ratios of the neurons **B** Same as in **A**, but when applied to motion-corrected recordings, with the exact same activity/firing as the static ones (see Methods) **C** Sorted accuracy levels for all spike sorters and recording types, as a function of the total number of neurons present in all artificial recordings **D** The run times for all the methods and datasets **E** For all spike sorting methods and recording types, the number of Well Detected, False Positive, Redundant and Overmerged units (see Methods).

When recordings have drift and are motion-corrected, there is a large decrease in performance for all spike sorters, as displayed in Figure 7B. As we saw in previous figures, this is mostly because both clustering and template-matching steps are severely impacted by the interpolation required when correcting for motion. Motion-correction degrades the overall number of cells that can be recovered, slows all pipelines (Figure 7D), and increases the number of False Positives, especially for Kilosort4 (see Figure 7E).

`Kilosort4` performs well on our generated data. It finds many good units and runs quickly, although is the only sorter that runs on a GPU. `Kilosort4` finds many false positives, especially for the motion-corrected recording. Importantly, this demonstrates that our simulated data poses a challenge to state-of-the-art spike sorters. The two spike sorters that were rebuilt as chains of modular components (`SpyKING-CIRCUS 2` and `TriDesClous 2`) have several pros and cons. `SpyKING-CIRCUS 2` has less False Positives but longer run times (Figure 7D, E). On the other hand, `TriDesClous 2` is fast, mostly due to its matching engine (working only on peak times). Such an implementation choice degrades its performances for static recordings but in the case of motion-corrected ones, the effect is less pronounced. However, the main advantages of these sorters lie in their modularity, so that it is straightforward to test ideas, customize pipelines and adapt parameters depending on the data. Finally, `Lupin` is designed to be the best combination of all currently available components on our dataset. It clearly outperforms all other sorters in all conditions (static or motion-corrected, including `Kilosort4`), with a good trade-off between the high number of found units, the speed, and the small number of False Positives. This is somewhat expected since such a pipeline is built with the “best” algorithms, evaluated on our generated data, carefully chosen at each step of the spike sorting pipeline. Its performance can be seen as a direct illustration of the gain obtained by sharing ideas within the community.

When designing `Lupin` we chose the component algorithms which performed best on our Neuropixel 1.0-like generated data. To demonstrate that the results shown in this paper generalize to new probe geometries, we performed additional simulations on ground truth datasets with Neuropixel 2.0, tetrode, SiNAPS [44] and Cambridge Neurotech layouts (see Supplementary Figure S1). In all cases, we found the same general trend observed in Figure 7, showing that `Lupin`’s success generalizes to other probe geometries. In all these cases, our modular pipeline `Lupin` was on par with state of the art sorters such as `Kilosort4` [18], with less False Positives and Redundant Units, especially in the presence of motion.

Finally, we tested the four spike sorters on real world datasets. We chose three datasets with diverse properties (chronic and acute, and from Cambridge Neurotech probes as well as Neuropixels 1.0 and 2.0 probes) and launched end-to-end spike sorters on the data. It is very difficult to assess spike sorters since there is no ground truth for any high-density ephys data. Hence we use results from three automatic curation algorithms as proxies to assess the sorting quality. Both Bombcell [45] and `UnitRefine` [46] label putative neurons as “good", “mua" (multi-unit activity) and “noise", providing approximate measures of “well detected", “overmerged" and “redundant" units. The `SLAy` [47] algorithm suggests units that should be merged, a proxy measure for “oversplit". Figure 8A shows the results of the automatic curation algorithms, with grouped columns corresponding to our categories. Note that these metrics are imperfect and only give a suggestion of the quality of the results. Similar to what has been observed with synthetic data, we can see that `Lupin` is competitive with `Kilosort4`, finding on average a similar number of good units and less noisy units. TriDesClous2 and SpyKINGCIRCUS 2 find fewer units, but have different advantages that might be interesting given the context. TriDesClous2 is fast, while SpyKING-CIRCUS 2 is mostly tailored for *in-vitro* data, and thus can scale for more thousands of channels while some other algorithms might not [18]. Overall the results show that our method, of optimizing and constructing a spike sorter based on carefully benchmarked results on simulated data, can generate a spike sorter on par with cutting edge end-to-end algorithms when applied to real data. As one can see in Figure 8B, which shows the activity over time for all recordings, the results do not seem to be related to the level of motion artifacts.

**Figure 8:**
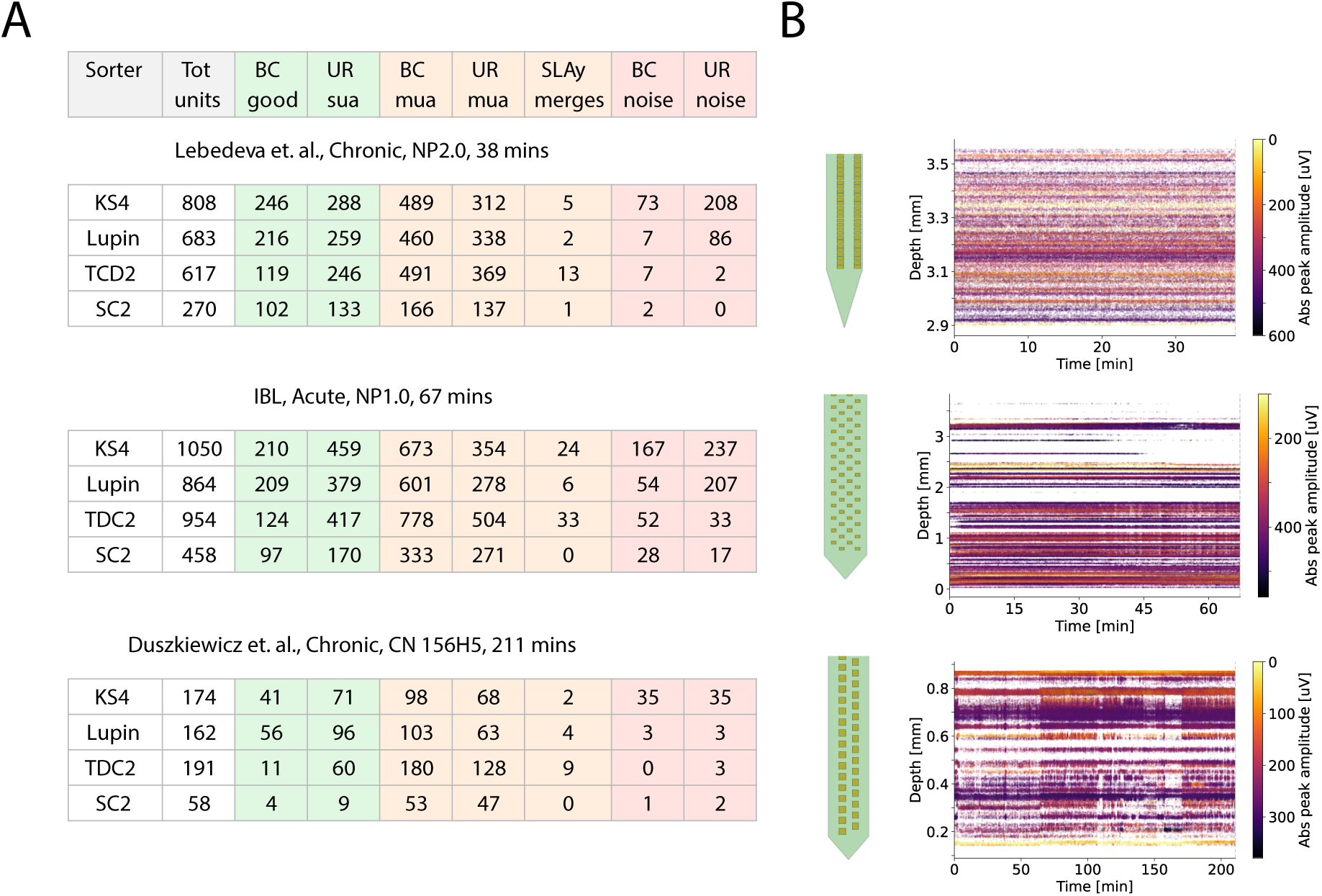
End-to-end spike sorter benchmarks on real data. Results from running four spike sorters (Kilosort4, Lupin, TriDesClous 2 and SpyKING-CIRCUS 2) on three datasets: a chronic Neuropixels 2.0 recording, an acute Neuropixels 1.0 recording [48, 49] and a long chronic recording using a 64-channel Cambridge Neurotech probe [50, 51] **A**: Tabulated proxy measurements of well-detected, overmerged or oversplit, and redundant units. For well-detected, the proxies were units being labeled “good” by `Bombcell` [45] or being labeled “sua” by the Jain lightweight model using `UnitRefine` [46]. The overmerged proxies were being labeled “mua” by `Bombcell` or the Jain model, and the oversplit proxy is the number of suggested merges from the `SLAy` algorithm [47]. The redundant proxies were being labeled “noise” by `Bombcell` or the Jain model. **B**: a representation of the probe used in each recording, and a raster map of spikes over time, giving an indication of the stability of each recording.

## Discussion

In this work, we have presented a new and modular framework to dissect and benchmark all steps of modern spike sorters, alongside a fast and efficient ground truth generator to quickly create artificial data to challenge the algorithms. This has allowed us to get a better understanding of the pros and cons of each individual algorithmic step, and allowed us to design new component-based spike sorters SpyKING-CIRCUS 2, TriDesClous 2 and Lupin. The latter sorter, Lupin, which is built using the best-performing method for each step, is already competitive with the most widely used sorter (Kilosort4) on our simulations and on real data. Unlike Kilosort4 our new sorters have been designed for high performance on CPUs, which is especially advantageous in the context of High Performance Computing and considering the cost and difficulty of getting access to GPUs.

We hope that our initial effort of dissecting and re-implementing several methods for each step of any spike sorter into a single unified framework will be beneficial for the spike sorting community at several levels. First, users now have access to a diverse range of adaptable spike sorters to obtain more accurate spike trains from their recordings. Second, developers of new methods will now be able to focus on a specific detail of a specific step (e.g., motion estimation, feature extraction, clustering, or template matching). They can implement new ideas without needing to write a completely new end-to-end sorter that includes the entire machinery (data reader, preprocessing, and user interface) since SpikeInterface already handles all these details. Finally, advanced users will be able to construct from scratch, or tweak, their own spike sorting solution that best fits their needs: a balance between accuracy, computational costs, and speed.

One could argue that such a modular based-approach might still have some limitations. First, it might appear as a suboptimal framework for spike sorters that run different steps iteratively rather than serially. Iterative methods have been used to refine some processing steps, e.g., having a re-clustering step after a first template-matching pass (e.g., as in Kilosort 1 and 2 [10], although note that Kilosort4 is a serial algorithm). However, nothing prevents creating iterative schemes and repeating individual steps in our modular framework, since all steps are implemented independently of the others. Second, our step-wise benchmarking effort could be extended and made more granular. For example, we did not benchmark the “feature extraction” step individually, but only in conjunction with clustering since most spike sorting algorithms extract spike features in a similar manner. But one could break the two steps up in order to properly benchmark each of them individually. This might provide an understand of how different feature extractions boost performance of particular clustering methods [26]. Third, this modular architecture might not reflect the complexity of “monolithic” spike sorting algorithms that have been optimized to work in specific manners. For example, failures at one particular step might be compensated for in subsequent steps in built-in pipelines, and these nonlinear optimizations might not be found in such a component-based framework. However, even if this is the case, identifying the sources of failures is beneficial to understand, and enhance the performances and the robustness of, the algorithms. The fact that `Lupin`, created here from scratch as the “optimal” combinations of components is *on par* with state of the art spike sorters both on artificial and real data (see Figures 7, 8, S1) is a direct demonstration that our optimization strategy works well.

We have demonstrated the validity of our component-based framework to benchmark individual methods and to build full end-to-end spike sorters, by using a fast and efficient way of generating ground truth data. Although the generative model that we use to simulate ground truth recordings does not capture all the complexities observed in experimental settings and could be enhanced, we believe that it already has the key ingredients to challenge spike sorting algorithms, such as cell inhomogeneities, randomness, and drift. It has the potential to speed up the development and comparison of spike sorters, not only for benchmarking purposes but also maybe, in the future, for the generation of training/labeled dataset that could be beneficial for end-to-end deep network solutions [52, 53].

In order to benchmark some of the individual steps, such as peak detection and clustering, datasets with partial ground truth (such as paired recordings or “hybrid” recordings) cannot be used (in peak detection, for example, the ground truth spikes will only be a very minor portion of all spikes in the recording). An alternative source of full ground truth datasets comes from biophysical simulations, which use advanced multi-compartment models to simulate extracellular templates [54] or even the full network activity that gives rise to extracellular recordings [55]. Although the latter types of simulation appear more biophysically plausible, it should be noted that virtually all available cell models are built from *in-vitro* experiments, which affect neuronal physiology and results in waveform shapes and distributions that are different from what is observed *in-vivo* [18]. A second drawback of biophysically plausible simulations is their computational complexity. Our proposed simulation framework trades off some biophysical “correctness” for efficiency and speed: our simulated data are generated almost instantaneously, on-the-fly and in memory upon request, requiring no disk space. Another idea that could be viable for some of the steps that do not require exhaustive ground truth (e.g. template matching or end-to-end comparisons) is hybrid recordings [18, 56]. A potential problem for hybrid recordings is one of circularity when it comes to motion correction. Hybrid data are built from experimental recordings by injecting ground truth spikes (from collections of templates or pre-curated spike sorting outputs). Since drift is a major phenomenon for shank-like probes, injected spikes should follow the inherent drift of the recordings. Spikes are therefore moved and interpolated given the estimated motion. When benchmarking the performance of spike sorters on such datasets, a spike sorter that uses the same motion correction method as is used to inject spikes will be favored. It is therefore important to make sure that motion correction is consistent across methods and potentially run prior to benchmarking.

While it has already been shown that drift in a recording causes a major degradation in spike sorting performance [19], in this work we further highlighted that the problem occurs most strongly in the clustering and template matching steps. It is mainly due to the way templates and/or traces are interpolated, even using state-of-the-art motion-correction algorithms [20, 57]. Template matching is very sensitive to residuals, and slight mismatches have a large impact on performance. The resolution of such issues remains an open challenge in spike sorting, and our work directly addresses this situation in the field: recordings with high drift amplitudes can have a very poor spike sorting quality. Therefore, it is important to stress that experimenters should try to minimize the drift during acquisition, since motion correction only works to a certain extent. In our opinion, if the estimated motion vectors are below 10 µm, it is better to turn off motion correction and work directly on the raw data. Such motion estimations can easily be computed and visualized with custom methods in `SpikeInterface`, to gain confidence in final results. One potential solution to improve motion correction is to make use of data from the new generation of ultra-dense probes, such as Neuropixels Ultra [58]. These novel probes provide an unprecedented spatial resolution, which could help to gain new insights on biophysical features of extracellular templates and be used to design better interpolation methods for the templates. It is also worth noting that switching from Neuropixels 1.0 to Neuropixels 2.0 seems to reduce the loss of well-detected units in motion corrected recordings compared to static recordings, regardless of the spike sorter (see Supplementary Figure S1 and Figure 7). This suggests that the higher density of electrodes (along the direction of the probe) found in Neuropixel 2 probes reduces the loss of accuracy due to motion.

Embedding this modular spike sorting framework in the SpikeInterface framework facilitates continued and distributed maintenance of the new modules. SpikeInterface is a mature ecosystem with several core developers across multiple institutes, over one hundred external contributors, an extensive testing suite and continuous integration across multiple operating systems and software infrastructure (e.g., Python versions). Finally, all the methods and options evaluated in this work are readily available to the electrophysiology community within SpikeInterface and can be immediately deployed with a few lines of code.

## Methods

### Notation

Throughout the article, vector variables are represented using⃗ notation and convolution using ∗ notation. We consider that the extracellular signals s⃗(t) are defined on N channels. We use w⃗_i_(t) ∈ ℝ^N×M^ to represent the spatio-temporal waveforms emitted by the neuron i at time t, where N is the total number of channels and M is the number of time samples. We further use the term Ground Truth (GT) to refer to the fully controlled variables in our synthetic recordings (either the motion signal or the spike times of the units).

### Data and code availability

All the figures available in this article can be regenerated from Jupyter notebooks that are available at https://zenodo.org/records/20407862 or https://github.com/SpikeInterface/sorting_components_benchmark_paper. All the methods and options evaluated in this work are readily available to the electrophysiology community within the SpikeInterface package https://github.com/SpikeInterface/spikeinterface and can be immediately deployed with a few lines of code. The implementations of the Kilosort clustering and matching are available at https://github.com/SpikeInterface/spikeinterface-kilosort-components.

### Simulated datasets

We have implemented a new `generation` module in `SpikeInterface` to ease the generation of ground truth artificial recordings, with or without motion drift to mimic *in-vivo* experiments [59]. This new strategy allows us to bypass the use of the `MEArec` simulator [54], at the cost of a slight loss in pseudo-realism. To create a simulated recording, a user must first provide a probe layout supported by `probeinterface` [60]. An in memory recording is created, which can produce traces on-the-fly, avoiding the need to save the generated recording to disk. This lazy mode is crucial, since ground truth recordings can be quite large for high-density probes with long durations. Given the templates, spike times, and the motion vector ^M⃗^(t) that affects the units, a seeded reproducible recording is created. Similar to what was done in the `MEArec` simulator, the user can control the number of units, their spike times and their physical positions in space. The spatio-temporal templates of the units w⃗(t) can either be extracted from third-party libraries or generated *via* a simple generative mathematical model.

To generate a template given the position of a neuron, we first generate a single prototypal waveform as a sum of decaying exponentials with various time constants to create the typical bi-phasic shape often observed *in-vivo*. This generation procedure has several parameters such as the peak negative/positive amplitudes, the time constant of depolarization and repolarization, and the recovery time. Parameters are randomly drawn from uniform distributions to make them different for every cell. Once a single waveform has been generated, it is scaled on every nearby channel by a spatial decay factor, formulated as a power law on the distances between the cell and channel positions. In order to ensure spatial anisotropies while computing these distances, we modeled the cell as an elongated ellipsoid whose axes and rotation are also randomized. Finally, the model takes into account a propagation speed such that waveforms are also temporally shifted as a function of the distances. Combined together we firmly believe that the model is able to reproduce the core features needed to properly challenge modern spike sorters, even while not capturing all the diversity observed *in-vivo* with respect to cell types, morphologies, etc. The precise implementation details can be found by inspecting the generation module of SpikeInterface.

A key feature of this new module is that it can handle motion drifts in two different ways. First, if cells are generated *via* the biophysical generative model, then they all have a source position p⃗ `= (`x, y, z) that can be varied as a function of motion ^M⃗^(t) during the course of the experiment. The positions (x, y) are randomly drawn within the area covered by the probe layout, extended by a 20 µm margin. Depths z are considered within the range [5, 40] µm. Second, if the unit templates are taken from an external library or provided by the user, the module will handle drift by interpolating these templates *via* cubic spline interpolation while shifting them with respect to the motion vector ^M⃗^(t).

The noise structure of these artificial recordings is simpler than observed in real data, but there are some key properties that can be specified by the user to challenge spike sorters. For example, the user can specify the noise levels per channel and even impose a given covariance matrix for the noise structure within the channels of the probes. Note, however, that the module does not consider any model of Local Field Potentials, thus making it only suitable for spike sorting benchmarks. Similar to [19], since the user has full access to the ground truth, the generation module can be used to impose a particular spatial distribution for the neurons (to mimic the layered organization of recorded structures), a particular distribution of firing rates and/or activity profiles (to mimic particular subtypes), and motion drifts that can be non-homogeneous as a function of the electrode depths.

In this paper, we focus mainly on 30 minute long simulated recordings with a Neuropixels-1.0 probe layout, generated with a sampling rate of 30 kHz. The number of units is fixed to 500, with a trimodal distribution of their positions along with the depth of the electrodes. Cell firing rates are drawn from a gamma distribution (shape 1 and scale 5) leading to rates from 0.1 to 30Hz. In Supplementary Figure S1, we generated four other simulated recordings each with a different probe layout but other conditions kept constant. Here, we use: a Cambridge NeuroTech ASSY-158-H5 64-channel layout; a Neuropixels 2.0-like 128 channel layout; a high density 128-channel SiNAPS layout; and a single 4-channel Tetrode. For all simulations, we fix the average cell density to 2.5 cells per channel.

### Real datasets

We tested the new components-based sorters on real data. To do so, we chose three datasets representing a diverse range of experimental conditions: a chronic implantation of a Neuropixels 2.0 probe from the Lebedeva et. al.^3^ [4, 61]; an acute implantation using a Neuropixels 1.0 probe from IBL^4^ [48, 49] and a long chronic recording using a Cambridge Neurotech probe from Duszkiewicz et. al.^5^ [50, 51]. We ran Lupin, SpyKING-CIRCUS 2 and TriDesClous 2, as well as Kilosort4 (version 4.1.2), on each dataset once without any additional preprocessing. The only default setting we adjusted was to apply motion correction or not depending on whether the recording was acute or chronic.

After sorting, we computed features of the spike sortings using `SpikeInterface`‘s sorting analyzer tools (including computing average waveforms, spike amplitude distributions, and template and quality metrics) allowing us to apply several automatic “quality control” tools each output. We applied `BombCell` [45], using default settings, which returns a list of units considered “sua” (singlue-unit activity), “mua” (multi-unit activity) and “noise” determined by a thresholding decision tree. We also applied the “lightweight” models from Jain using `UnitRefine` [46] These models are classifiers trained on thousands of manually curated units from several datasets. The models output the quality labels “sua”, “mua” and “noise”, and should represent the decisions of the manual curation decisions fed to the models. Finally, we applied `SLAy` an auto-merging tool [47], which returns paired units it determines to be originally oversplit by the sorters.

Since there is no ground truth for real spike sorted data, we use the output of these tools as a proxy for the categories we assessed the simulated data on: well-detected, oversplit and redundant. We use the `Bombcell` “sua" labels and the `UnitRefine` “sua” label as proxies for well-detected units, `Bombcell` “noise” and `UnitRefine` “noise” labels as proxies for redundant units, `Bombcell` “mua” and `UnitRefine` “mua” labels as a proxies for overmerged units, and number of suggested `SLAy` merges a proxy for oversplit units. These are imperfect metrics. For instance, both the default `Bombcell` parameter tuning the Jain model [46] use Neuropixels data sorted using `Kilosort4` as their input. Regardless, we felt it was fairest to use the default settings to judge the sorting outputs.

### Peak detection

#### Locally exclusive

In this method, commonly performed in spike sorters [15, 11, 17], peaks are detected as negative threshold crossings within a spatio-temporal exclusion zone, to avoid cross-contamination between channels. This is implemented as the “locally exclusive” method of `SpikeInterface`. The parameters of such a detection method are the detection threshold, defined as how many times above the median absolute deviation that the signal must be to be counted as a peak (`detection_threshold = 5`), the spatial radius in which such a peak must be a local extrema (`local_radius_um = 50` µm) and the temporal exclusion zone during which such an extrema must be unique (`exclude_sweep_ms = 2ms`).

#### Matched filtering

This peak detection method relies on the the concept of matched filtering [37]. Because spikes have stereotypical temporal waveforms, we can increase the signal-to-noise ratio of the peaks during the detection process by specifically looking for such shapes in the signal. Introduced in [18], the idea behind this peak detection method is to create an exhaustive catalog F of artificial templates ^f⃗^_n∈1,…k_(t) at known positions p⃗_n_ `= (`x_n_, y_n_), and to convolve the extracellular signal s⃗(t) with all these spatio-temporal templates to find matches in the data.

To create these artificial templates, the typical waveform H⃗(t) of some detected peaks on a single channel is estimated as a median of, say, `5`, `000` normalized waveforms. These initial waveforms are taken from peaks that are detected using the locally exclusive method described above. The computed single channel waveform is duplicated on all nearby channels, such that on every channel c at a position p⃗_c_ `= (`x_c_, y_c_) we have ^f⃗^_n_(c, t`) =` w_n_(c)H⃗(t) with a weight w_n_(c) that will reflect the spatial decay of the templates. In the following, we model the spatial decay as:

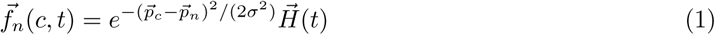

In this formula σ controls the spatial decay of the templates and to further extend the catalog, multiple values of σ can be used (in the range of 10 to 50 µm). Once the catalog is obtained, all these templates F are convolved with the extracellular signal s⃗(t), giving rise to a new signal r⃗(t) ∈ ℝ^K^, where K is the number of templates. Typically, K is much larger than N, the number of channels.

The final peaks are then detected via the aforementioned locally exclusive method, but on this new signal r⃗(t). Note that because the stereotypical waveform H⃗(t) is the same as the one used to generate all the spatio-temporal templates in the catalog F, the convolutions between F and s⃗(t) can be highly optimized by only performing the convolution between s⃗(t) and H⃗(t) per channel, and then summing these individual convolutions with the weights w_n_(c).

### Motion estimation

The motion estimation and correction steps of the spike sorting pipeline were modularized in a previous paper [19]. Although SpikeInterface offers multiple methods to handle motion correction, we believe that there is sufficient evidence that DREDge provides the best motion estimation [62, 20]. Hence throughout the paper we always use DREDge to infer the motion of our recordings.

Since motion correction is now a canonical step in most preprocessing pipelines for *in-vivo* recordings with high-density probes, it is important to assess how well various algorithms are with respect to this key preprocessing step. Hence, we often compare the results obtained on static recordings with “motioncorrected” recordings. To be precise, for every ground-truth recording that has been generated using the `generation` module, we create both a static and a drifting version of the recording. A “motioncorrected” recording is a drifting recording in which the motion has been estimated by `DREDge` and then compensated for *via* a kriging interpolation [18]. To our knowledge, this is the closest, albeit not perfect, way to estimate the static recording from the drifting one. If this step was perfect, the “motion-corrected" recording would be equal to the “static" one. Even if such a motion estimation step is pretty good [19], it might still distorts the signal in two ways. First, since it depends on activity levels and cell positions, we need to have many active cells in a localized region to properly estimate the motion [19]. So when there is little activity, the motion estimation is imperfect. Second, because the kriging interpolation used to compensate for the motion smooths the signal and noise, some information from the data is lost during this step.

### Clustering

Clustering is one of the most important steps in spike sorting, and numerous clustering methods have been developed. A complete review of all these methods is outside the scope of this manuscript. In the SpikeInterface components module, we have implemented some of the key modern approaches used for high-density probes. Most of these methods start from the observation that clustering in a high-dimensional space is an intractable problem, so one must reduce the dimensionality before clustering. The most obvious way to do so, while keeping the spatio-temporal features of the waveforms needed for clustering, is to perform a Singular Value Decomposition (SVD). To do so, several methods first gather single-channel waveforms w⃗_i_(t) ∈ ℝ^1×M^, where M is the number of time steps (usually, the temporal duration of a spike *in-vivo* lasts 2 ms), and then perform an SVD to reduce the number of dimensions from M to K `= 5`, with K < M. Assuming that such an SVD projection is rather typical, it can be extended on a per-channel basis to spatio-temporal waveforms such that snippets w⃗_i_(t) ∈ ℝ^N×M^ observed on N channels can be projected into a space ℝ^N×K^. In such lower-dimensional spaces, the clustering is easier. In the following, we use w⃗^SV^ ^D^ ∈ ℝ^N×K^ for the projected representation of w⃗_i_ ∈ ℝ^N×M^, after SVD. In all the following, we used K `= 5`, as is commonly used by many sorters.

#### KS-clustering

The full details of this clustering method can be found in [18]. In a “divide-and-conquer" approach, the waveforms w⃗^SV^ ^D^ are split into groups as a function of their estimated depth (the algorithm being tailored for Neuropixels-like probes), and a graph-based clustering algorithm is applied per group (as a modified version of the Louvain or Leiden algorithms [63]). The results of all these individual clusterings are then concatenated while ensuring that cells at the borders of the bin depths, which could give rise to duplicated clusters, are merged before template matching (duplicated templates are removed).

#### Iterative clusterings (Iter-HDBSCAN and Iter-ISOSPLIT)

These methods are similar to the one used in `Kilosort`, but rely on density-based clustering algorithms. In another “divide-and-conquer" approach, the waveforms w⃗^SV^ ^D^ are split into groups as a function of their estimated depth (the algorithm being tailored for Neuropixels-like probes), and a clustering algorithm is applied locally. In particular, all detected peaks are grouped with respect to their peak channel. For all these peaks, all channels in a given neighborhood are selected, and projected onto a low-dimensional space to obtain the w⃗^SV^ ^D^. These are then clustered, using a density-based clustering algorithm (such as HDBSCAN [64]) or one based on statistical considerations (such as ISOSPLIT [12, 65]). The results of all these individual clusterings are then concatenated while ensuring that cells at the border of the bin depths, which could give rise to duplicated clusters, are merged afterwards. To do this, these clustering algorithms contain an additional cleaning step that ensures: 1) all similarities measurements between found templates, as defined by an l1 norm, are less than a certain threshold (0.8) 2) All small clusters that are likely to be noisy are removed (given a firing rate threshold for found neurons).

#### Full graph sparse clustering (Global Louvain)

This method attempts to avoid the edge effect of the “divide-and-conquer" approaches encountered with iterative splits and local clusterings. The method also begins with projected waveforms w⃗^SV^ ^D^ however, unlike the previous methods, the graph-based clustering is applied to the full connectivity graph at once, in order to avoid border effects. A key to making this work is that distances (and thus edges) of the connectivity graph between all waveforms are only computed for waveforms that are spatially close enough. Once the connectivity matrix is computed for all local distances, we can apply the clustering algorithm of our choice (Louvain [63], or even HDBSCAN for the distances). In this manuscript, we used the Louvain algorithm.

### Template matching

Once the templates T⃗_i_(t) ∈ ℝ^N×M^ have been found and identified as the centroids of the clusters, template matching attempts to describe the signal s⃗(t) as a linear sum of the templates, plus noise. Here again, various methods have been tried, with small differences and subtleties.

#### KS-matching

This is a simple Matching Pursuit algorithm [66], and full details can be found in [18, 10]. The signal s⃗(t) is convolved with all the templates T⃗_i_(t) ∈ ℝ^N×M^. This convolution leads to the computation of all the scalar products b_ij_ `=` T⃗_i_(t_j_) · s⃗(t_j_) at any time t_j_. To speed up these convolutions, one can use an SVD representation of the templates, and/or optimized libraries such as `torch`. Once the b_ij_ are computed, the algorithm iteratively takes the one with the largest value b_i_∗_,j_∗ (i.e. the best match), and subtracts b_i_∗_,j_∗ T_i_∗ (t_j_∗ ) from the signal s⃗(t), leaving behind the residual. This procedure is repeated until no more matches can be found, leading, in theory and if the dictionary is complete, to residuals that should only represent the noise present in the signal s⃗(t). In practice, however, the dictionary of templates is not fully accurate, and a stopping criteria must be imposed. Here again, to speed up the algorithms, operations are performed in the space of the scalar products instead of the raw data. This requires precomputation of all pairwise scalar products of all pairs of templates T_i,j_(t) for all possible lags. Assuming that we have N_T_ templates T_i_(t) ∈ ℝ^N×M^, this lookup table M has a size T × T × `(2`M − `1)`. While not the case in `Kilosort`, such a lookup table can be sparsified in practice because not all templates interact with each other.

#### Circus-OMP

This is based on the Orthogonal Matching Pursuit algorithm [42], which is slightly different to the matching algorithm implemented in `Kilosort`. The main difference is that each time a template is selected (and thus b_i∗,j∗_ is chosen), this selection updates the values of b_i∗,j∗_that were selected so far. Intuitively, this means that adding a new template to the reconstruction updates the weights of the ones that were selected before. Such an algorithm has been shown to be more efficient when templates are non-orthogonal [67]. Again, to speed up the implementation, some internal optimizations can be performed, via Cholesky decomposition [68].

#### Wobble

This is an augmented Matching Pursuit algorithm, as implemented in YASS [17]. The ideas are similar to the classical Matching Pursuit Algorithm of `Kilosort`, but involve a super-resolution step to enhance the alignment of the templates. In summary, the dictionary of templates T⃗_i_(t) ∈ ℝ^N×M^ are “augmented” through multiple slightly time-delayed versions of the templates, to compensate for subsampling jittering. This produces a dictionary of T⃗ (t) ∈ ℝ^N^ ^×M^, with N^′^ > N, and then uses an algorithm similar to the one of Kilosort.

#### TDC-peeler

This is a greedy Matching Pursuit algorithm that works only at peak times, originally developed both in SpyKING-CIRCUS [15] and T. In short, peaks are detected as thresholds that cross above k times the median absolute deviation per channel. As in the peak detection step, a spatiotemporal exclusion radius ensures that only the most prominent peaks are kept. At these peak times, we look for the best match to the templates with respect to Euclidean distances. Once a match is found, it is subtracted from the raw traces, and peaks are redetected on the residuals traces until there are no further matches.

#### TDC-peeler drift aware

This is a special case of the “TDC-peeler" algorithm, which highlights the effect of motion correction while matching templates. It is not part of any of the components based sorters, and is only used in Figure 6. Instead of interpolating the traces, as is done by most modern sorters [18], we extend the dictionary of templates by moving them from −`100` µm to `100` µm along the y−axis *via* `1` µm spatial increments. Interpolation is performed with bi-cubic splines, and once this augmented set of templates has been generated, we apply exactly the same matching algorithm as “TDC-peeler", but now selecting the appropriate template at each time point, based on the inferred motion at this time point.

To generate Figure 5E, we used the same methodology as in [39] to assess how well the matching strategies were able to detect collisions. In particular, we used the extension of the ground-truth comparison class `CollisionGTComparison`, which computes performance metrics by spike lag. In addition to the agreement score computation and the matching, this method first detects and flags all “synchronous spike events” in the ground-truth spike trains. Two spikes from two separate units are considered a “synchronous spike event” if their spike times occur within a time delay of `2 ms`. The synchronous events are then divided into 11 bins that span the [−`2`, `2] ms` interval, and *collision recall* is computed for each bin. The similarities between templates are computed as the normalized l2 norm, with a number between 0 and 1 quantifying how similar two templates are. Collision recalls as function of the lags are plotted by grouping pairs of templates as a function of their similarities, with the assumption that the task of solving temporal collision is different if templates are dissimilar or not.

### End-to-end spike sorter comparison on synthetic recordings

We include here a brief description of all the end-to-end spike sorters compared in Figure 7: SpyKING-CIRCUS 2 This is an updated version of SpyKING-CIRCUS [15] based on the modular components implemented in this article. In summary, this spike sorter uses (when motion is present) the DREDge motion correction algorithm [20] before whitening the data. On this whitened data, the chain of components that are used are: matched filtering for peak detection, iterative splits for clustering (Iter-HDBSCAN), and orthogonal matching pursuit for template reconstruction (Circus-OMP). Finally, units are merged greedily with built-in function of SpikeInterface suggesting putative merges. The results presented in this paper were created using version 25.12.

TriDesClous 2 This is an updated version of TriDesClous based on the modular components implemented in this article. In summary, the code uses (when motion is present) the DREDge motion correction algorithm [20] before filtering the data. On this filtered data, the chain of components that are used are: locally exclusive for peak detection, iterative splits for clustering (Iter-ISOPLIT), and fast greedy partial deconvolution, only applied at peak times for template reconstruction (TDC-peeler). Finally, units are merged greedily with built-in function of SpikeInterface suggesting putative merges. The results presented in this paper were created using version 25.12.

Lupin This is a direct demonstration of the potential unlocked by the modular components implemented in this article. In summary, the code uses (when motion is present) the DREDge motion correction algorithm [20] before filtering and whitening the data. On this whitened data, the chain of components that are used are: matched filtering for peak detection, iterative splits for clustering (Iter-ISOPLIT), and augmented matching pursuit for the spike deconvolution (Wobble). Finally, units are merged greedily with built-in function of SpikeInterface suggesting putative merges. The results presented in this paper were created using version 25.12.

Kilosort4 This is the complete standalone algorithm, as implemented in [18]. All the units found by Kilosort were kept for downstream analysis. We used version 4.1.1.

### Evaluation

The exact nature of the evaluations that have been performed in every benchmark depends on the nature of the benchmark. In all of our benchmarks, we tried to identify the key goals of every step (for example, capture accurately all peaks for peak detection, find all clusters for clustering, etc.) and design quality metrics accordingly. Quite often, this relies on a comparison between the ground truth labels of the spike trains and the one provided as output by the different algorithms (in clustering, matching, merging). This comparison is performed based on the agreement matrix and the so-called “matches”. More details can be found in the `SpikeInterface` documentation but roughly, such matches allow us to compute the average accuracy per matched units. For some particular figures (such as clustering, matching, etc.), we also looked at the number of good, overmerged, and false positive units (see [34] for more details).

Knowing the ground-truth spiking activity, we can compute the *accuracy* of each ground-truth unit i as:

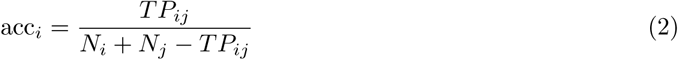

where j is the sorted unit matched to the Ground Truth (GT) unit i, N_i_and N_j_ are the number of spikes in the GT and the matched sorted unit, respectively, and T P_ij_ is the number of *true positive* spikes, i.e., the spikes found both in the GT and sorted spike trains. From this accuracy metric, we further classified spike sorted units as:

- *well detected* : units with an accuracy greater than or equal to 80%.
- *overmerged* : units with an agreement greater than 20% with more than one GT unit.
- *redundant* : units with an agreement above 20% with a single GT unit. These units can be oversplit or duplicated sorted units.
- *false positive*: sorted units with an agreement below 20%.

### Hardware Specifications

All simulations and spike sorting jobs have been run on a Intel(R) Xeon(R) Silver 4210 CPU @ 2.20GHz Machine with a Nvidia Quadro RTX 4000 GPU (used only by Kilosort).

## Acknowledgments

This work has been funded through several grants. We thank Joe Ziminski for his helpful feedback and comments on the manuscript. PY is supported by the ANR-24-RRII-000, the ANR GNEURO ANR-25-CE42-6535-03 and the *Cross-Disciplinary Project LOOP “Closed-Loop Neurotechnologies: from sensors to applications”* (Initiative d’excellence Université de Lille R-CDP-25-003-LOOP. ZM is supported by NIH grants F31129103, T32GM007753, T32GM144273. CH is supported by the UKRI Biotechnology and Biological Sciences Research Council (BBSRC) grant number BB/X01861X/1. PAF, HRM, and BD received support by NIH grant U19NS123716-02. CF is supported by NSF Neuronex Award DBI-1707398 and Simons Foundation grant 344 543023. We further thank the Simons Foundation for supporting the SpikeInterface project.

## Supplementary Figures

**Figure S1:**
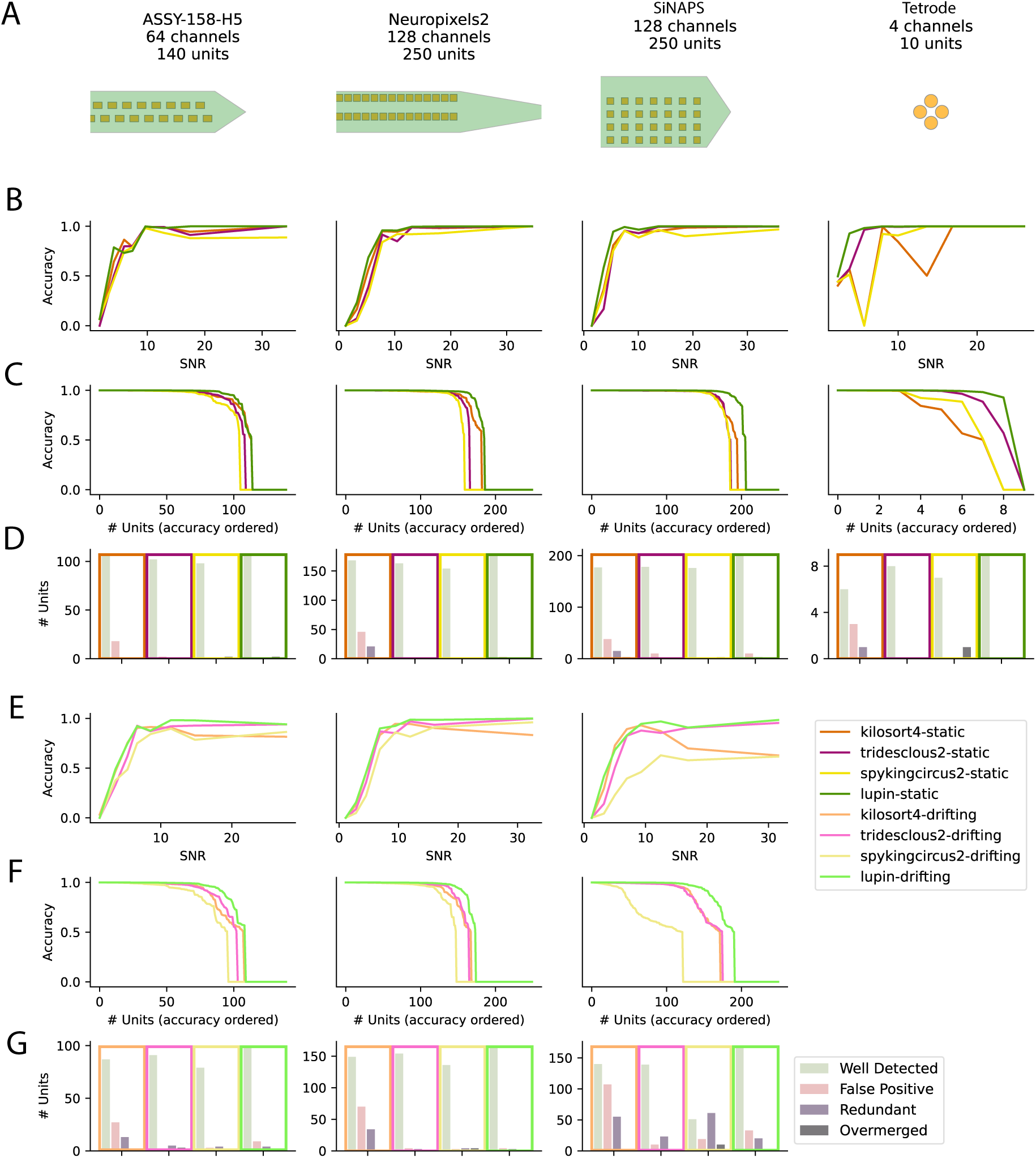
End-to-end spike sorter benchmark with extra simulated recordings. **A** Different probe geometries considered to generate the synthetic ground truth (see Methods). **B** Averaged accuracy for several spike sorters when applied to several static recordings, as a function of the signal-to-noise ratios of the neurons **C** Sorted accuracy levels for all spike sorters and recording types, as a function of the total number of neurons present in all artificial recordings **D** For all spike sorting methods and recording types, the number of Well Detected, False Positive, Redundant and Overmerged units (see Methods) **E, F, G** Same as in **B, C, D**, but when applied to motion-corrected recordings, with the exact same activity/firing as the static ones (see Methods).

## Footnotes

1 While many of these sorters are integrated in the SpikeInterface framework, their original GitHub repositories have had no commits over the last two years

2 used with a detection threshold of 5 median absolute deviations, which is “standard” in spike sorting [38]

3 Using recording AL032_2020-01-07

4 Using recording sub-UCLA034_ses-3537d970-f515-4786-853f-23de525e110f_desc-raw_ecephys.nwb

5 Using recording sub-A3702_ses-191126_behavior+ecephys.nwb

## References

[1] A. P. Buccino, S. Garcia, and P. Yger, “Spike sorting: new trends and challenges of the era of high-density probes,” Progress in Biomedical Engineering, vol. 4, no. 2, p. 022005, 2022.

[2] B. Lefebvre, P. Yger, and O. Marre, “Recent progress in multi-electrode spike sorting methods,” Journal of Physiology-Paris, vol. 110, no. 4, pp. 327–335, 2016.

[3] J. J. Jun, N. A. Steinmetz, J. H. Siegle, D. J. Denman, M. Bauza, B. Barbarits, A. K. Lee, C. A. Anastassiou, A. Andrei, Ç. Aydın, et al., “Fully integrated silicon probes for high-density recording of neural activity,” Nature, vol. 551, no. 7679, p. 232, 2017.

[4] N. A. Steinmetz, C. Aydin, A. Lebedeva, M. Okun, M. Pachitariu, M. Bauza, M. Beau, J. Bhagat, C. Böhm, M. Broux, et al., “Neuropixels 2.0: A miniaturized high-density probe for stable, longterm brain recordings,” Science, vol. 372, no. 6539, 2021.

[5] G. N. Angotzi, F. Boi, A. Lecomte, E. Miele, M. Malerba, S. Zucca, A. Casile, and L. Berdondini, “Sinaps: An implantable active pixel sensor cmos-probe for simultaneous large-scale neural recordings,” Biosensors and Bioelectronics, vol. 126, pp. 355–364, 2019.

[6] S. Hua, Y. Liu, J. Luo, S. Li, L. Jiang, P. Wu, S. Sun, L. Shang, C. Lu, K. Zhang, J. Liu, M. Wang, H. Shi, and X. Cai, “Microelectrode arrays cultured with in vitro neural networks for motion control tasks: encoding and decoding progress and advances,” Microsystems & Nanoengineering, vol. 11, no. 1, p. 233, 2025.

[7] R. Wolff, A. Polito, A. P. Buccino, M. Chiappalone, and V. Tucci, “Protocol for the enhanced analysis of electrophysiological data from high-density multi-electrode arrays with nicespike and spikeNburst,” STAR protocols, vol. 6, no. 4, p. 104195, 2025.

[8] M. Schröter, F. Cardes, C.-V. H. Bui, L. D. Dodi, T. Gänswein, J. Bartram, L. Sadiraj, P. Hornauer, S. Kumar, M. Pascual-Garcia, et al., “Advances in large-scale electrophysiology with highdensity microelectrode arrays,” Lab on a Chip, vol. 25, no. 19, pp. 4844–4885, 2025.

[9] C. Rossant, S. N. Kadir, D. F. Goodman, J. Schulman, M. L. Hunter, A. B. Saleem, A. Grosmark, M. Belluscio, G. H. Denfield, A. S. Ecker, et al., “Spike sorting for large, dense electrode arrays,” Nature neuroscience, vol. 19, no. 4, p. 634, 2016.

[10] M. Pachitariu, N. A. Steinmetz, S. N. Kadir, et al., “Fast and accurate spike sorting of high-channel count probes with kilosort,” in Advances in Neural Information Processing Systems, pp. 4448–4456, 2016.

[11] G. Hilgen, M. Sorbaro, S. Pirmoradian, J.-O. Muthmann, I. E. Kepiro, S. Ullo, C. J. Ramirez, A. P. Encinas, A. Maccione, L. Berdondini, et al., “Unsupervised spike sorting for large-scale, high-density multielectrode arrays,” Cell reports, vol. 18, no. 10, pp. 2521–2532, 2017.

[12] J. E. Chung, J. F. Magland, A. H. Barnett, et al., “A fully automated approach to spike sorting,” Neuron, vol. 95, no. 6, pp. 1381–1394, 2017.

[13] J. J. Jun, C. Mitelut, C. Lai, S. L. Gratiy, C. A. Anastassiou, and T. D. Harris, “Real-time spike sorting platform for high-density extracellular probes with ground-truth validation and drift correction,” BioRxiv, p. 101030, 2017.

[14] R. Diggelmann, M. Fiscella, A. Hierlemann, and F. Franke, “Automatic spike sorting for highdensity microelectrode arrays,” Journal of neurophysiology, vol. 120, no. 6, pp. 3155–3171, 2018.

[15] P. Yger, G. L. Spampinato, E. Esposito, B. Lefebvre, S. Deny, C. Gardella, M. Stimberg, F. Jetter, G. Zeck, S. Picaud, et al., “A spike sorting toolbox for up to thousands of electrodes validated with ground truth recordings in vitro and in vivo,” Elife, vol. 7, p. e34518, 2018.

[16] F. J. Chaure, H. G. Rey, and R. Quian Quiroga, “A novel and fully automatic spike-sorting implementation with variable number of features,” Journal of neurophysiology, vol. 120, no. 4, pp. 1859–1871, 2018.

[17] J. Lee, C. Mitelut, H. Shokri, I. Kinsella, N. Dethe, S. Wu, K. Li, E. B. Reyes, D. Turcu, E. Batty, et al., “Yass: Yet another spike sorter applied to large-scale multi-electrode array recordings in primate retina,” bioRxiv, 2020.

[18] M. Pachitariu, S. Sridhar, J. Pennington, and C. Stringer, “Spike sorting with Kilosort4,” Nature Methods, vol. 21, pp. 914–921, May 2024. Publisher: Nature Publishing Group.

[19] S. Garcia, C. Windolf, J. Boussard, B. Dichter, A. P. Buccino, and P. Yger, “A Modular Implementation to Handle and Benchmark Drift Correction for High-Density Extracellular Recordings,” eNeuro, vol. 11, pp. ENEURO.0229–23.2023, Feb. 2024.

[20] C. Windolf, H. Yu, A. C. Paulk, D. Meszéna, W. Muñoz, J. Boussard, R. Hardstone, I. Caprara, M. Jamali, Y. Kfir, D. Xu, J. E. Chung, K. K. Sellers, Z. Ye, J. Shaker, A. Lebedeva, R. T. Raghavan, E. Trautmann, M. Melin, J. Couto, S. Garcia, B. Coughlin, M. Elmaleh, D. Christianson, J. D. W. Greenlee, C. Horváth, R. Fiáth, I. Ulbert, M. A. Long, J. A. Movshon, M. N. Shadlen, M. M. Churchland, A. K. Churchland, N. A. Steinmetz, E. F. Chang, J. S. Schweitzer, Z. M. Williams, S. S. Cash, L. Paninski, and E. Varol, “DREDge: robust motion correction for high-density extracellular recordings across species,” Nature Methods, vol. 22, pp. 788–800, Apr. 2025.

[21] K. J. Laboy-Juárez, S. Ahn, and D. E. Feldman, “A normalized template matching method for improving spike detection in extracellular voltage recordings,” Scientific reports, vol. 9, no. 1, p. 12087, 2019.

[22] G. Saggese, M. Tambaro, E. A. Vallicelli, A. G. Strollo, S. Vassanelli, A. Baschirotto, and M. D. Matteis, “Comparison of sneo-based neural spike detection algorithms for implantable multitransistor array biosensors,” Electronics, vol. 10, no. 4, p. 410, 2021.

[23] R. Toosi, M. A. Akhaee, and M.-R. A. Dehaqani, “An automatic spike sorting algorithm based on adaptive spike detection and a mixture of skew-t distributions,” Scientific reports, vol. 11, no. 1, p. 13925, 2021.

[24] B. C. Souza, V. Lopes-dos Santos, J. Bacelo, and A. B. Tort, “Spike sorting with gaussian mixture models,” Scientific reports, vol. 9, no. 1, p. 3627, 2019.

[25] J. Eom, I. Y. Park, S. Kim, H. Jang, S. Park, Y. Huh, and D. Hwang, “Deep-learned spike representations and sorting via an ensemble of auto-encoders,” Neural Networks, vol. 134, pp. 131– 142, 2021.

[26] J. Wouters, F. Kloosterman, and A. Bertrand, “A data-driven spike sorting feature map for resolving spike overlap in the feature space,” Journal of Neural Engineering, vol. 18, no. 4, p. 0460a7, 2021.

[27] Y. Zhang, J. Han, T. Liu, Z. Yang, W. Chen, and S. Zhang, “A robust spike sorting method based on the joint optimization of linear discrimination analysis and density peaks,” Scientific reports, vol. 12, no. 1, p. 15504, 2022.

[28] M. R. Keshtkaran and Z. Yang, “Noise-robust unsupervised spike sorting based on discriminative subspace learning with outlier handling,” Journal of neural engineering, vol. 14, no. 3, p. 036003, 2017.

[29] P. S. Shabestari, A. P. Buccino, S. S. Kumar, A. Pedrocchi, and A. Hierlemann, “A modulated template-matching approach to improve spike sorting of bursting neurons,” in 2021 IEEE Biomedical Circuits and Systems Conference (BioCAS), pp. 1–4, IEEE, 2021.

[30] J. Wouters, F. Kloosterman, and A. Bertrand, “Towards online spike sorting for high-density neural probes using discriminative template matching with suppression of interfering spikes,” Journal of neural engineering, vol. 15, no. 5, p. 056005, 2018.

[31] J. Wouters, F. Kloosterman, and A. Bertrand, “A data-driven regularization approach for template matching in spike sorting with high-density neural probes,” in 2019 41st Annual International Conference of the IEEE Engineering in Medicine and Biology Society (EMBC), pp. 4376–4379, IEEE, 2019.

[32] J. Magland, J. J. Jun, E. Lovero, A. J. Morley, C. L. Hurwitz, A. P. Buccino, S. Garcia, and A. H. Barnett, “Spikeforest, reproducible web-facing ground-truth validation of automated neural spike sorters,” Elife, vol. 9, p. e55167, 2020.

[33] C. Merow, B. Boyle, B. J. Enquist, X. Feng, J. M. Kass, B. S. Maitner, B. McGill, H. Owens, D. S. Park, A. Paz, et al., “Better incentives are needed to reward academic software development,” Nature Ecology & Evolution, vol. 7, no. 5, pp. 626–627, 2023.

[34] A. P. Buccino, C. L. Hurwitz, S. Garcia, J. Magland, J. H. Siegle, R. Hurwitz, and M. H. Hennig, “Spikeinterface, a unified framework for spike sorting,” Elife, vol. 9, p. e61834, 2020.

[35] A. J. G. Wyngaard, V. Llobet, and B. Barbour, “Lussac: a fully-automated consensus method that increases the yield and quality of spike-sorting analyses.” Pages: 2022.02.08.479192 Section: New Results.

[36] P. N. Steinmetz, “Simulation of background neuronal activity and noise in human intracranial microwire recordings,” Journal of Neuroscience Methods, vol. 402, p. 110017, 2024.

[37] G. Turin, “An introduction to matched filters,” IRE Transactions on Information Theory, vol. 6, no. 3, pp. 311–329, 1960.

[38] R. Q. Quiroga, Z. Nadasdy, and Y. Ben-Shaul, “Unsupervised spike detection and sorting with wavelets and superparamagnetic clustering.,” Neural computation, vol. 16, no. 8, pp. 1661–1687, 2004. ISBN: 0899-7667 (Print).

[39] S. Garcia, A. P. Buccino, and P. Yger, “How do spike collisions affect spike sorting performance?,” Eneuro, vol. 9, no. 5, 2022.

[40] M. Pachitariu, N. A. Steinmetz, and J. Colonell, “Kilosort2,” 2019. https://github.com/MouseLand/Kilosort2.

[41] M. Pachitariu, S. Sridhar, and C. Stringer, “Solving the spike sorting problem with kilosort,” bioRxiv, pp. 2023–01, 2023.

[42] Y. Pati, R. Rezaiifar, and P. Krishnaprasad, “Orthogonal matching pursuit: recursive function approximation with applications to wavelet decomposition,” in Proceedings of 27th Asilomar Conference on Signals, Systems and Computers, pp. 40–44 vol.1, Nov. 1993. ISSN: 1058-6393.

[43] J. Boussard, C. Windolf, C. Hurwitz, H. D. Lee, H. Yu, O. Winter, and L. Paninski, “DARTsort: A modular drift tracking spike sorter for high-density multi-electrode probes,” Aug. 2023. Pages: 2023.08.11.553023 Section: New Results.

[44] G. N. Angotzi, M. Vöröslakos, N. Perentos, J. F. Ribeiro, M. Vincenzi, F. Boi, A. Lecomte, G. Orban, A. Genewsky, G. Schwesig, D. Aykan, G. Buzsáki, A. Sirota, and L. Berdondini, “Multi-shank 1024 channels active SiNAPS probe for large multi-regional topographical electrophysiological mapping of neural dynamics,” Advanced Science, vol. 12, no. 16, p. 2416239, 2025.

[45] J. M. J. Fabre, E. H. v. Beest, A. J. Peters, M. Carandini, and K. D. Harris, “Bombcell: automated curation and cell classification of spike-sorted electrophysiology data.”

[46] A. Jain, R. Greene, C. Halcrow, J. A. Swann, A. Kleinjohann, F. Spurio, S. Graff, A. Pan-Vazquez, B. Kampa, J. Gall, S. Grün, O. Winter, A. Buccino, M. H. Hennig, and S. Musall, “UnitRefine: A community toolbox for automated spike sorting curation.”

[47] S. Koukuntla, T. DeWeese, A. Cheng, R. Mildren, A. Lawrence, A. R. Graves, K. E. Cullen, J. Colonell, T. D. Harris, and A. S. Charles, “SLAy-ing oversplitting errors in high-density electrophysiology spike sorting,” bioRxiv: The Preprint Server for Biology, p. 2025.06.20.660590, 2025.

[48] D. Angelaki, B. Benson, J. Benson, D. Birman, N. Bonacchi, K. Bougrova, S. A. Bruijns, M. Carandini, J. A. Catarino, et al., “A brain-wide map of neural activity during complex behaviour,” Nature, vol. 645, no. 8079, pp. 177–191, 2025.

[49] International Brain Laboratory, B. Benson, J. Benson, D. Birman, N. Bonacchi, M. Carandini, J. Catarino, G. Chapuis, P. Dayan, E. DeWitt, T. Engel, M. Fabbri, M. Faulkner, I. Fiete, C. Findling, L. Freitas-Silva, B. Gerçek, K. Harris, S. Hofer, F. Hu, F. Hubert, J. Huntenburg, A. Khanal, C. Langdon, P. Lau, G. Meijer, N. Miska, J.-P. Noel, K. Nylund, A. Pan-Vazquez, A. Pouget, C. Rossant, N. Roth, R. Schaeffer, M. Schartner, Y. Shi, K. Socha, N. Steinmetz, K. Svoboda, A. Urai, M. Wells, S. West, M. Whiteway, O. Winter, and I. Witten, “Ibl - brain wide map.” DANDI:DANDI:000409/0.260309.1324.

[50] A. J. Duszkiewicz, P. Orhan, S. Skromne Carrasco, E. H. Brown, E. Owczarek, G. R. Vite, E. R. Wood, and A. Peyrache, “Local origin of excitatory–inhibitory tuning equivalence in a cortical network,” Nature neuroscience, vol. 27, no. 4, pp. 782–792, 2024.

[51] A. Duszkiewicz, S. Skromne Carrasco, and A. Peyrache, “Large-scale recordings of head direction cells in mouse postsubiculum.” DANDI:DANDI:000939/0.250207.0025.

[52] Y. He, A. Marin-Llobet, H. Sheng, R. Liu, and J. Liu, “End-to-end multimodal deep learning for real-time decoding of months-long neural activity from the same cells,” Oct. 2024. Pages: 2024.10.14.618046 Section: New Results.

[53] Y. Han and S. Wang, “E-Sort: Empowering End-to-end Neural Network for Multi-channel Spike Sorting with Transfer Learning and Fast Post-processing,” Dec. 2024. arXiv:2409.13067 [eess].

[54] A. P. Buccino and G. T. Einevoll, “Mearec: a fast and customizable testbench simulator for ground-truth extracellular spiking activity,” Neuroinformatics, pp. 1–20, 2020.

[55] S. Laquitaine, M. Imbeni, J. Tharayil, J. B. Isbister, and M. W. Reimann, “Spike sorting biases and information loss in a detailed cortical model,” BioRxiv, pp. 2024–12, 2024.

[56] A. P. Buccino, A. Sridhar, D. Feng, K. Svoboda, and J. H. Siegle, “Efficient and reproducible pipelines for spike sorting large-scale electrophysiology data,” Elife, Feb. 2026.

[57] N. Watters, A. Buccino, and M. Jazayeri, “MEDiCINe: Motion correction for neural electrophysiology recordings.” Pages: 2024.11.06.622160 Section: New Results.

[58] Z. Ye, A. M. Shelton, J. R. Shaker, J. Boussard, J. Colonell, D. Birman, S. Manavi, S. Chen, C. Windolf, C. Hurwitz, H. Yu, T. Namima, F. Pedraja, S. Weiss, B. C. Raducanu, T. V. Ness, X. Jia, G. Mastroberardino, L. F. Rossi, M. Carandini, M. Häusser, G. T. Einevoll, G. Laurent, N. B. Sawtell, W. Bair, A. Pasupathy, C. M. Lopez, B. Dutta, L. Paninski, J. H. Siegle, C. Koch, S. R. Olsen, T. D. Harris, and N. A. Steinmetz, “Ultra-high-density Neuropixels probes improve detection and identification in neuronal recordings,” Neuron, vol. 113, pp. 3966–3982.e12, Dec. 2025. Publisher: Elsevier.

[59] N. Steinmetz, “"imposed motion datasets" from steinmetz et al. science 2021,” Feb 2021.

[60] S. Garcia, J. Sprenger, T. Holtzman, and A. P. Buccino, “ProbeInterface: A Unified Framework for Probe Handling in Extracellular Electrophysiology,” Frontiers in Neuroinformatics, vol. 16, p. 823056, Feb. 2022.

[61] A. Lebedeva, M. Okun, M. K. Krumins, and M. Carandini, “Chronic recordings from Neuropixels 2.0 probes in mice,” 12 2023.

[62] E. Varol, J. Boussard, N. Dethe, O. Winter, A. Urai, T. I. B. Laboratory, A. Churchland, N. Steinmetz, and L. Paninski, “Decentralized motion inference and registration of neuropixel data,” in ICASSP 2021-2021 IEEE International Conference on Acoustics, Speech and Signal Processing (ICASSP), pp. 1085–1089, IEEE, 2021.

[63] V. D. Blondel, J.-L. Guillaume, R. Lambiotte, and E. Lefebvre, “Fast unfolding of communities in large networks,” Journal of Statistical Mechanics: Theory and Experiment, vol. 2008, p. P10008, Oct. 2008.

[64] L. McInnes, J. Healy, and S. Astels, “hdbscan: Hierarchical density based clustering,” Journal of Open Source Software, vol. 2, p. 205, Mar. 2017.

[65] J. F. Magland and A. H. Barnett, “Unimodal clustering using isotonic regression: ISO-SPLIT,” May 2016. arXiv:1508.04841 [stat].

[66] S. Mallat and Z. Zhang, “Matching pursuits with time-frequency dictionaries,” IEEE Transactions on Signal Processing, vol. 41, pp. 3397–3415, Dec. 1993.

[67] S. Bian and L. Zhang, “Overview of match pursuit algorithms and application comparison in image reconstruction,” in 2021 IEEE Asia-Pacific Conference on Image Processing, Electronics and Computers (IPEC), pp. 216–221, 2021.

[68] A. Lubonja, S. K. Præsius, and T. D. Tran, “Efficient Batched CPU/GPU Implementation of Orthogonal Matching Pursuit for Python,” July 2024. arXiv:2407.06434 [cs].

